# Analysis and extension of exact mean-field theory with dynamic synaptic currents

**DOI:** 10.1101/2021.09.01.458563

**Authors:** Giulio Ruffini

## Abstract

Neural mass models such as the Jansen-Rit or Wendling systems provide a practical framework for representing and interpreting electrophysiological activity (1–6) in both local and global brain models (7, 8). However, they are only partly derived from first principles. While the post-synaptic potential dynamics are inferred from data and can be grounded on diffusion physics (9–11), Freeman’s “wave to pulse” (W2P) sigmoid function (12–14), used to transduce mean population membrane potential into firing rate, rests on a weaker theoretical standing. On the other hand, Montbrió et al (15, 16) derive an exact mean-field theory (MPR) from a quadratic integrate and fire neuron model under some simplifying assumptions, thereby connecting microscale neural mechanisms and meso/macroscopic phenomena. The MPR model can be seen to replace Freeman’s W2P sigmoid function with a pair of differential equations for the mean membrane potential and firing rate variables—a dynamical relation between firing rate and membrane potential—, providing a more fundamental interpretation of the semi-empirical NMM sigmoid parameters. In doing so, we show it sheds light on the mechanisms behind enhanced network response to weak but uniform perturbations. In the exact mean-field theory, intrinsic population connectivity modulates the steady-state firing rate W2P relation in a monotonic manner, with increasing self-connectivity leading to higher firing rates. This provides a plausible mechanism for the enhanced response of densely connected networks to weak, uniform inputs such as the electric fields produced by non-invasive brain stimulation. This new, dynamic W2P relation also endows the neural mass model with a form of “inertia”, an intrinsic delay to external inputs that depends on, e.g., self-coupling strength and state of the system. Next, we complete the MPR model by adding the second-order equations for delayed post-synaptic currents and the coupling term with an external electric field, bringing together the MPR and the usual NMM formalisms into a unified exact mean-field theory (NMM2) displaying rich dynamical features. In the single population model, we show that the resonant sensitivity to weak alternating electric field is enhanced by increased self-connectivity and slow synapses.

**Significance**
Several decades of research suggest that weak electric fields influence neural processing. A long-standing question in the field is how networks of neurons process spatially uniform weak inputs that barely affect a single neuron but that produce measurable effects in networks. Answering this can help implement electric field coupling mechanisms in neural mass models of the whole brain, and better represent the impact of electrical stimulation or ephaptic communication. This issue can be studied using local detailed computational models, but the use of statistical mechanics methods can deliver “mean-field models” to simplify the analysis. Following the steps of Montbrió et al (15, 16), we show that the sensitivity to inputs such a weak alternating electric field can be modulated by the intrinsic self-connectivity of a neural population, and produce a more grounded set of equations for neural mass modeling to guide further work.

## 1. Introduction

In their seminal paper, Montbrió et al (15) provide a rigorous derivation of population dynamics from first principles using statistical mechanics methods. Their framework (MPR) provides a derivation of the neuron equations of state relating membrane potential and firing rate, in contrast to the prescription of a sigmoid function representation of the firing rate-membrane potential relation (W2P function) as is done in earlier work. This allows to properly interpret the meaning and limitations of the effective theory. In particular, it can help define effective electrical interaction models of neural masses with weak external fields. The new W2P function is dynamic, and as a result endows the model with new properties, such as delayed response to inputs.

In their derivation, Montbrió et al made several assumptions, some by necessity and others for simplicity. In particular, they use instantaneous synaptic currents, which makes the model easier to analyze but also unrealistic and hard to relate to neural mass modeling, where synaptic delays play a key role. This has been recently expanded to include first-order synapse equations in (17, 18). We complement here the MPR equations with second-order post-synaptic kinetics equations as used in NMM (4, 9–11, 19) to represent post-synaptic potentials (PSPs). By bringing these theories together, a new, extended model emerges (NMM2) providing a strong link between microscopic mechanisms and parameters and the mesoscopic population phenomena described by semi-empirical neural mass models.

In the next sections, we summarize the NMM formalism and analyze its linearized version to show that the effective coupling to external inputs is modulated by several terms, including connectivity and sigmoid baseline rate. We then review the MPR model, add to it dimensional constants, and extend it to include second-order synapse dynamics, multiple interacting populations (NMM2 model) and interaction with a uniform external electric field. We show (v. SI section A) that the interaction is of dipole form if the neuron is far from threshold (in a linear regime or the field is very weak).

We then show that the single population model W2P function, which replaces the usual sigmoid in NMM, is a function of self-connectivity or quenched noise parameters (excitability), and that changing either modulates the sensitivity of the mass to external inputs and modulates the resonant frequency and amplitude to weak perturbations. Moreover, the dynamics of the new firing rate-membrane potential relation (W2P function) lead to a delayed response of the mass to external inputs (which we call mass “inertia”), with the delay a function of self-connectivity. We show that a self-inhibitory model can oscillate. Finally, we produce a two population model (Excitation-Inhibition or E-I), and present some of its dynamics, which are far richer than the classical analog neural mass model.

### Neural mass models

Neural mass (semi-empirical) models (NMM), first developed in the early seventies by W. Freeman (20) and F. Lopez de Silva (2), provide a physiologically grounded description of the average synaptic activity and firing rate of a neural population (1–5, 8). NMMs are increasingly used for local and whole-brain modeling in neurology (e.g., epilepsy (6, 21) or Alzheimer’s disease (22)) and for understanding and optimizing the effects of transcranial electrical brain stimulation (tES) (7, 8, 23, 24). The central conceptual elements in this framework are the synapse, which is seen to transduce incoming activity (quantified by firing rate) into a mean membrane potential perturbation in the receiving neuron population, and the sigmoid function transforming population membrane potential to output (mean) firing rate with due account for threshold and saturation effects (see (19) for a nice introduction to the Jansen-Rit model). The synaptic filter is instantiated by a second-order system coupling the mean firing rate of arriving signals *φ* to the mean post-synaptic voltage perturbation *u*,

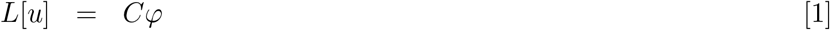

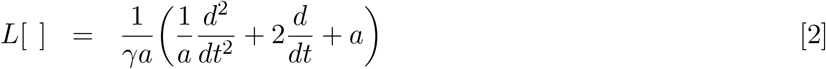

or, in the equivalent first-order form,

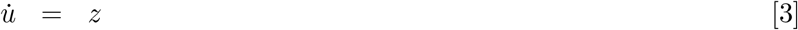

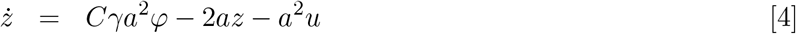

where *u*(*t*) is the mean post-synaptic voltage perturbation produced by the synapse, *φ*(*t*) is the mean input firing rate and *C* is the dimensionless connectivity constant quantifying the mean number of synapses per neuron in the receiving population. The action of the *L* operator can also be represented by its inverse, a convolution with its Green’s function *h*(*t*) = *H*(*t*)*γa*^2^*te*^− *at*^ (with *H*(*t*) the Heaviside step function) (19).

These linear ordinary differential equations can be generalized to account for different rising and decay times (9, 11). The parameters *a* and *γ* describe the delay time scale 1*/a* in seconds and the amplification factor in V/Hz (traditionally (19), the parameter *A* (V) is used instead of *γ*, with *A* = *γa*). The parameter *C* is dimensionless and quantifies the average number of synapses per neuron in the receiving population. The synaptic transmission equations need to be complemented by a relationship between the membrane potential of the neuron and its firing rate (the neuron W2P function). Freeman proposed the sigmoid as a simple model capturing essential properties of the response of neuron populations from inputs (12–14), based on modeling insights and empirical observations. Note this is a static relation between membrane potential and firing rate. Each neuronal population 𝒫_*n*_ converts the sum *v*_*n*_ of the membrane perturbations from each of the incoming synapses or external perturbations (e.g., an external electric field) to an output firing rate (*φ*_*n*_) in a non-linear manner using Freeman’s sigmoid function,

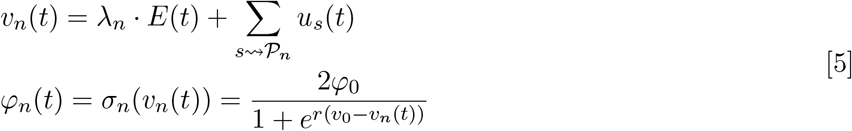

where *φ*_0_ is half of the maximum firing rate of each neuronal population, *v*_0_ is the value of the potential when the firing rate is *φ*_0_ and *r* determines the slope of the sigmoid at the central symmetry point (*v*_0_, *φ*_0_) (we have omitted an *n* subscript in these population dependent parameters for simplicity). The sum is over synapses *onto* population 𝒫_*n*_, which we express as 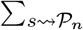. The effect of an electric field is represented by a membrane potential perturbation, *δv*_*n*_ = *λ*_*n*_·*E*, with *λ*_*n*_ an effective, semi-empirical vector parameter that represents the first-order dipole coupling of field and neuron.

The presence of semi-empirical “lumped” quantities such as *φ*_0_ and *λ* highlight the fact that NMMs are not currently derived from microscopic models, but bring together a mix of theory and observational relations. This means, for example, that even if we have access to a detailed single neuron compartment model providing an electric dipole coupling constant, it is not obvious how to use it in an NMM. To better understand the effective coupling to external inputs, we consider a small perturbation of the population mean membrane potential around an equilibrium point 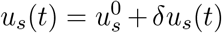, that is, 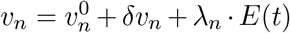 with *λ*_*n*_ · *E*(*t*) a weak field perturbation and 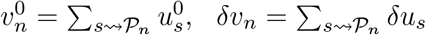. Then, we can linearize the sigmoid around the fixed point (Taylor expansion to first order) to obtain

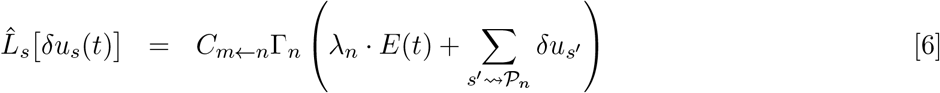

where *C*_*m*←*n*_ is connectivity matrix from population *n* to *m* and 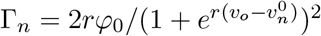. In the regime of small perturbations around an equilibrium point, the effective electric coupling impact is given by the product of *λ*_*n*_ *C*_*m*←*n*_ Γ_*n*_. However, it is not obvious how the parameters involved should be changed as a function of the neural mass properties (e.g, cortical path size it represents or self-connectivity), because in NMM they are semi-empirical quantities.

## 2. Extended MPR framework (NMM2)

### MPR model

In their uniform, mean-field derivation for a population of *N* quadratic integrate and fire (QIF) neurons, Monbrió et al (15) start from the equations for the neuron membrane potential perturbation from baseline,

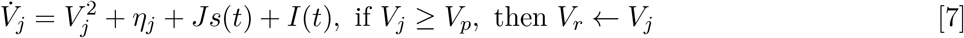

In this equation, the total input current in neuron *j* is *I*_*j*_ = *η*_*j*_ + *Js*(*t*) + *I*(*t*) and includes a quenched noise constant component *η*_*j*_, the input from other neurons *s*(*t*) per connection received (the mean synaptic activation) with uniform coupling *J* and a common input *I*(*t*). The common input *I*(*t*) can represent both a common external input or the effect of an electric field, i.e.,

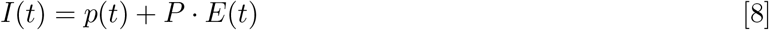

Here *p*(*t*) is an external uniform current, and *P* is the dipole conductance term in the spherical harmonic expansion of the response of the neuron to an external, uniform electric field. This is a good approximation if the neuron is in its subthreshold, linear regime (see SI section A), and can be computed using realistic compartment models of the (see, e.g., (25) and (26)).

The mean synaptic activation is given by

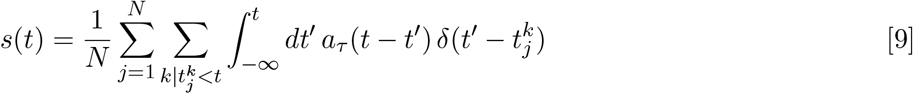

where 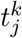 is the arrival time of the *k*th spike from the *j*th neuron, and *a*(*t*) the synaptic activation function, e.g., *a*(*t*) = *e*^− *t/τ*^ */τ*. Note that we can write *J* = *j·N*, where *j* is the synapse coupling strength (charge delivered to the neuron per action potential at the synapse) of each synapse the cell receives from the network (there are *N* of them in a fully connected architecture with *N* neurons).

We assume here for simplicity that all neurons are equally oriented with respect to the electric field. If the electric field is constant, variations in orientation can be absorbed by the quenched noise term. The total input *p*(*t*) + *P* · *E*(*t*) + *Js*(*t*) is thus homogeneous across the population (does not depend on the neuron).

Starting from these, Montbrió et al derive an effective theory in the large *N* limit (Eq 12 in (15)),

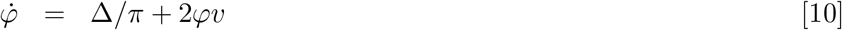

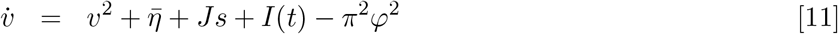

Here *v* and *φ* are the population mean membrane potential and firing rate respectively. The new parameters 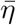 and Δ refer to the mean and half-width of the quenched noise distribution of *η*_*j*_. The analysis in (15) hinges on the assumptions of all to all uniform connectivity (with synaptic weight *J*) and common input *I*(*t*). In addition, a set of closed equations can be derived on the limit of instantaneous synaptic transmission (*τ* → 0 or *a*_*τ*_(*t*− *t*′) →*δ*(*t* − *t*′)), which implies *s* → *φ* to produce a closed, simple set of equations for the single population model,

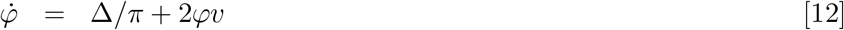

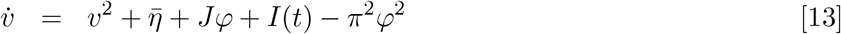

This zero delay model is the one analyzed in (15). Figure 1 provides a diagram of the self-coupled population.

**Fig. 1.**
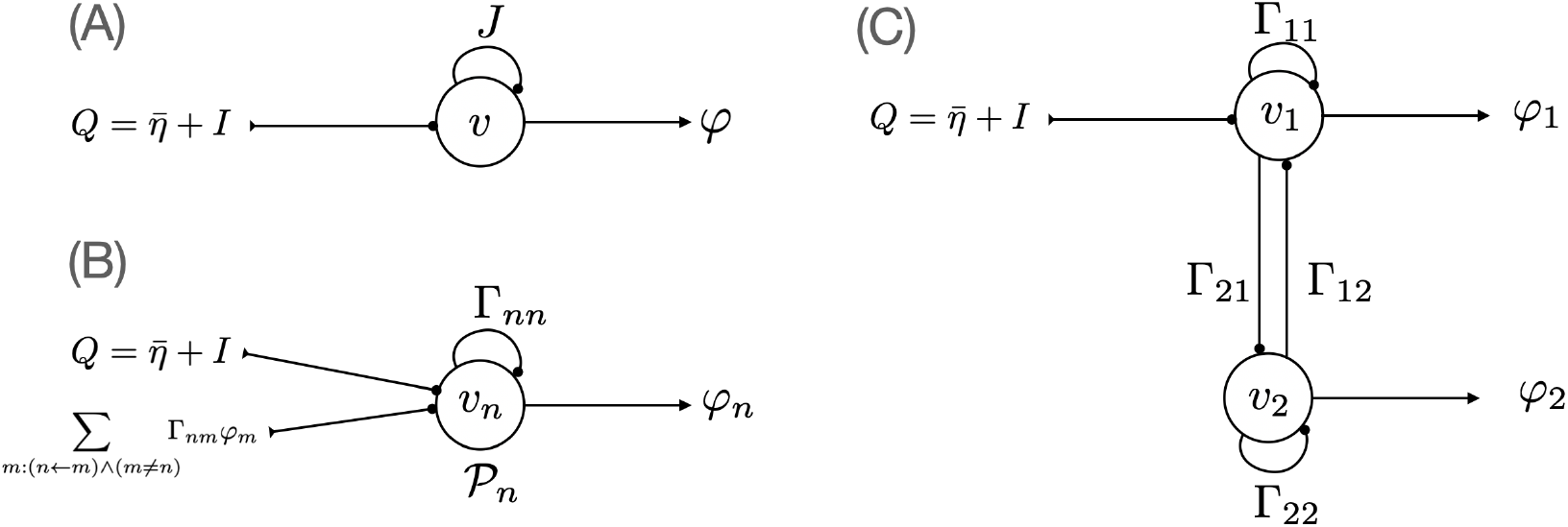
NMM2 diagrams. (A): Diagram for self-coupled population with connectivity *J* receiving and external input *Q*. (B): generalization for multiple populations. (C): A generic two population model.

**Fig. 2.**
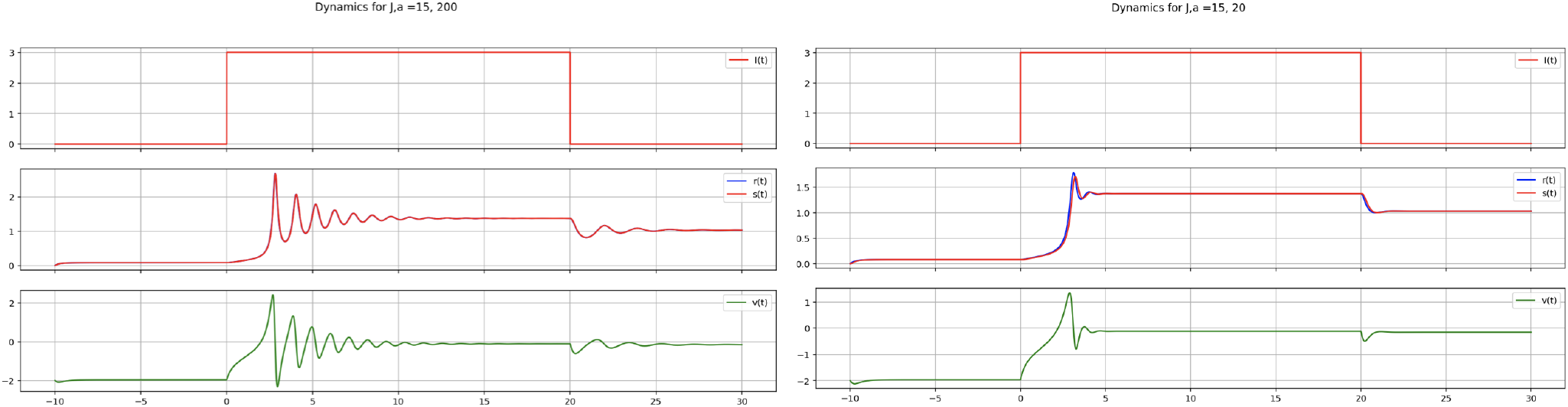
Dynamics of NMM2 single population model. Left: Fast synaptic dynamics (with rate constant *a* = 200) are equivalent to the original MPR model where *s* = *φ*. Right: Slower dynamics (*a* = 20) with effect of synaptic filtering (*J* = 15, *η* = − 5, Δ = 1).

### Adding dimensional constants

In order to dimensionalize the MPR model to make connection with experimental work, we start from the full QIF formulation in (17),

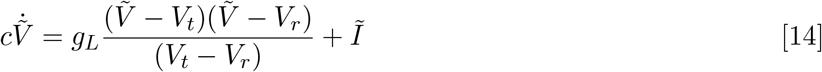

where *V*_*r*_ and *V*_*t*_ represent the resting potential and threshold of the neuron, 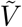 is the membrane voltage, *Ĩ* the input current, *c* the membrane capacitance, *g*_*L*_ is the leak conductance, with units 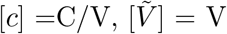,[*Ĩ*] = A and [*g*_*L*_] = A/V. To simplify this (completing the square), we define

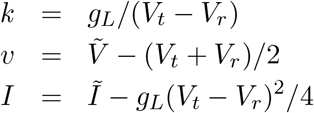

to obtain

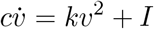

with *k* units [*k*]= A/V^2^ and voltage and current in proper units. Our starting point thus becomes the simplified QIF equation,

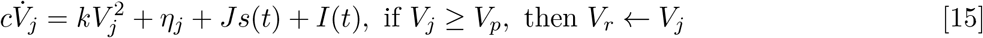

Following the derivation of the mean-field equations in (15) from this QIF formulation now leads to

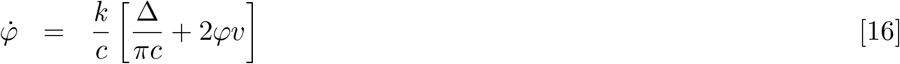

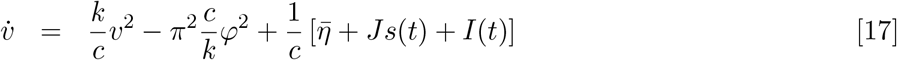

with units [*r, s*]=Hz, [*J*]=C, [*v*]=V, [*k*]=A/V^2^, [*c*]=C/V=A/(V Hz), 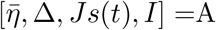, and [*k/c*]=Hz/V.

### Extended equations

Montbrió et al mention the use of an exponential decay function to relate firing rate and synaptic activity but for simplicity set *s* = *φ* in their final derivation. We extend this analysis here to make a connection with the neural mass modeling formalism.

When an action potential wave reaches the synapse, it is transformed into the influx of some ionic species into the cell. This transforms the shape of the action potential signal, low-pass filtering it. This process is governed by the Poisson-Nernst-Planck equation (see, e.g., (10)), but can be described in a simplified manner by the dynamics of conductance *g*(*t*) through the underlying ohmic relation *I*_*s*_ = *g*(*t*) (*V* − *V*_*R*_). As with the membrane perturbation, the dynamics of conductance can be modeled by a second-order linear operator (2) or its generalization to different rise and decay times (9, 11). To include synaptic dynamics we use a convolution operator as in (15), but redefine the synaptic activation function to include a Heaviside function (implementing causality), *h*_*τ*_(*t*) = *H*(*t*)*a*_*τ*_ (*t*). Then we can write (the convolution operator is linear)

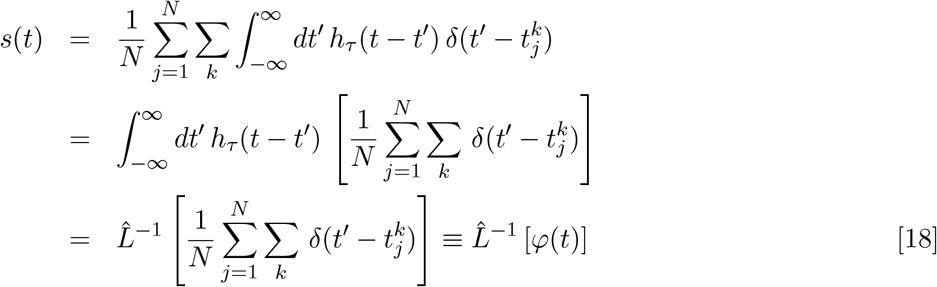

with 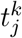 the arrival time of the *k*th spike from the *j*th neuron, and *φ*(*t*) the instantaneous firing rate. Thus, we have 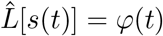 to account for input gate delay. We will use the second-order operator discussed in Equation 2 to represent post-synaptic current kinetics (the generalized version with different rise and decay times can be equally used). The full set of equations for a single population becomes

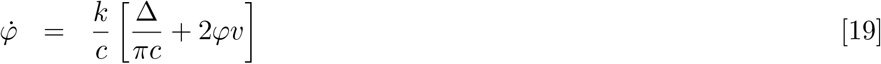

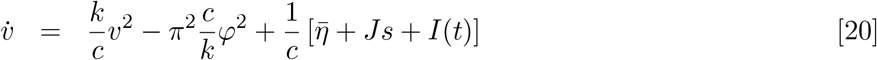

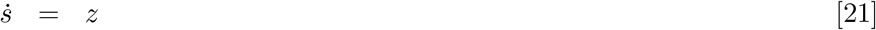

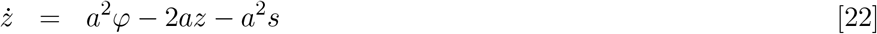

with the input collecting external input and the influence of an external field, *I*(*t*) = *p*(*t*) + *P* · *E*(*t*) (see SI section A and O). These equations can be reduced to a simplified form using dimensionless parameters (see SI section I),

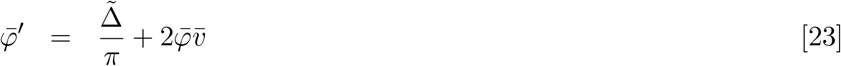

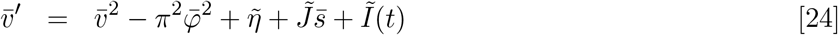

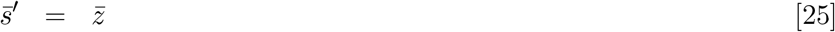

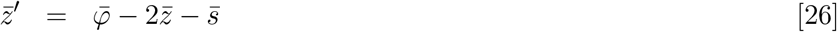

### Multiple population equations

The single population case is readily generalized for interacting populations. Letting *n* denote the population index and *nm* denote a synapse from population *m* to *n*, the uniform input received by a population is

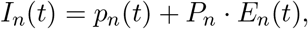

The equations for interacting populations become

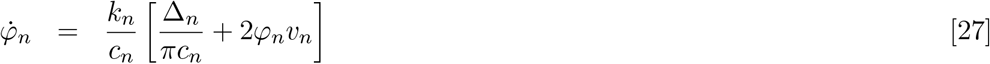

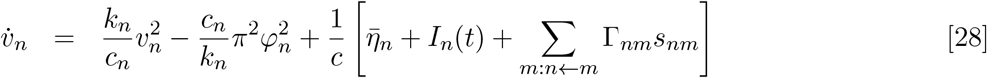

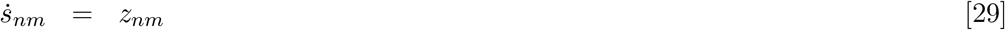

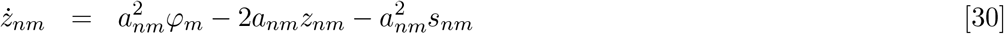

This can be further simplified by assuming that the synapses associated to a source neuron are all equivalent, then *s*_*nm*_ = *s*_*m*_ and *a*_*nm*_ = *a*_*m*_. The fixed points of these equations are independent of *a*_*nm*_ and are essentially the same for the “unextended” equations

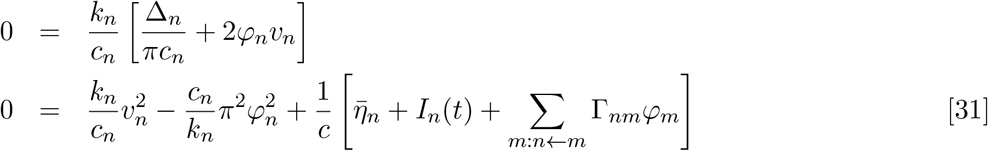

with *z*_*nm*_ = 0 and *s*_*nm*_ = *φ*_*m*_. The stability properties of these fixed points do depend on *a*_*nm*_, see, e.g., Figure 4 or Figure SI-12 and others in SI.

## 3. Analysis of single population NMM2 model

We consider here a single self-coupled population with constant input *I* and in steady state. Let *Q* represent the total input, 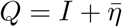. Then, the fixed point equations become

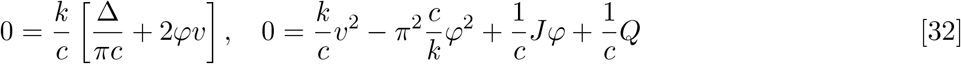

Substituting the first equation into the second we get

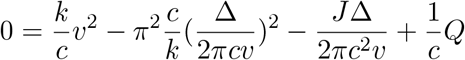

Canceling factors and multiplying by *v*^2^ we end with

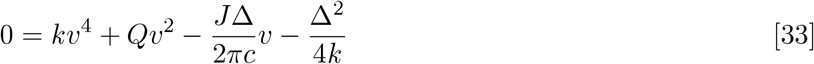

together with the supralinear neuron W2P function

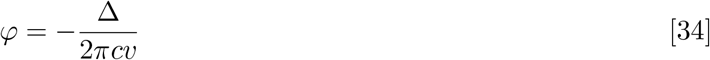

which requires *v* < 0 since *φ* ≥ 0. Equation 34 is a direct relation between voltage and firing rate and replaces the sigmoid in NMMs. Differentiating Equation 33 w.r.t. *J*, and using Equation 33 in the last term in the denominator, we find (since *v* < 0)

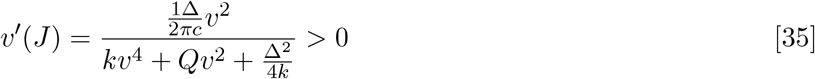

since *k* > 0. This in turn implies, using Equation 34,

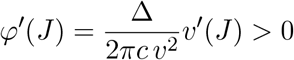

that is, membrane potential and firing rate monotonically increase with intrinsic connectivity, all other parameters being equal. Figure 3 displays the relation between firing rate and membrane potential as a function of connectivity *J* for a set of inputs. As can be observed, intrinsic connectivity modulates the operating point of the neuronal W2P function.

**Fig. 3.**
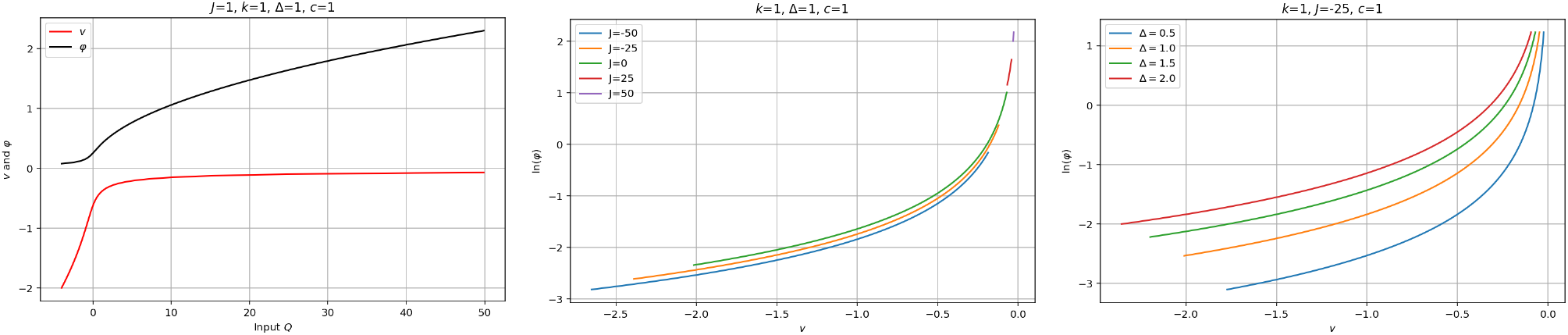
W2P function of single population model NMM2 as a function of connectivity *J* and Δ. Left: sample response to varying total input for a specific value of *J*. We observe the initial sigmoidal response shape of the firing rate *φ*. Middle: Relation of mean firing rate and membrane potential for a set of total input values *Q* ∈ (− 4, 50), as a function of intrinsic connectivity *J* (different color traces). The underlying neuronal W2P remains the same, but changing *J* moves the working point. NB: the traces for each value of *J* are sequentially slightly displaced upward for clarity, but they all lie on the same line, Equation 34. Right: same but as a function of quenched noise half-width Δ (without shifting the traces).

In summary, increasing connectivity leads to an increase of firing rate and membrane potential, all other things being equal. Thus, the intrinsic connectivity *J* acts like a (nonlinear but monotonic) gain modulator and affects the sensitivity of the population to external inputs. A *J* increases, so does the response of the system to an input. Translating this into the NMM framework means the sigmoid parameters such as *φ*_0_ (Hz) in NMMs should be adjusted to reflect intrinsic connectivity. To wit, *J*, the signed (negative if the population is inhibitory) average number of synapses per neuron in the population, is a scaling factor of the firing rate. As a consequence of the above, the sensitivity of dynamics to input depends greatly on the self-connectivity strength of each neural mass. Figure 4 displays this relationship. The impact on dynamics of this change in operating point of the W2P function will depend, in general, on the state of the system.

**Fig. 4.**
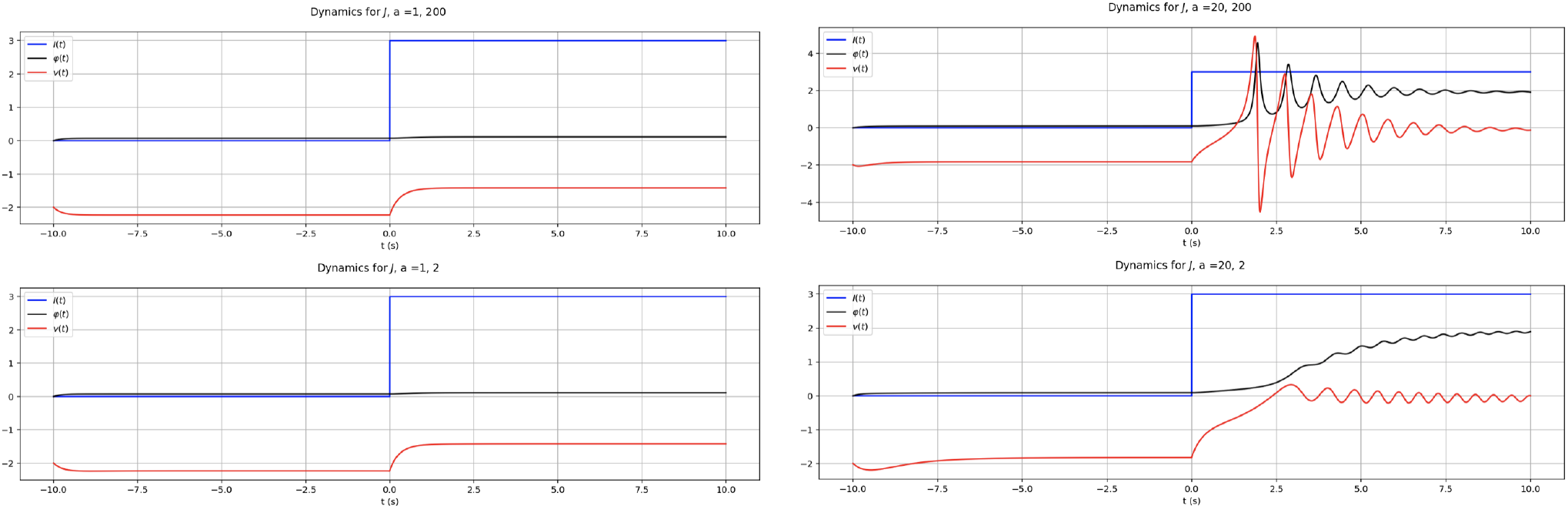
Sensitivity to input depends on *J* and rate constant in single population NMM2 model. Left: response to input with *J* = 1 for fast (top) and slow (bottom) synaptic dynamics. Right: response with stronger self-connectivity *J* = 20 for fast (top) and slow (bottom) synaptic dynamics. In both cases *η* =− 5, Δ = 1. See Figures SI-4 and SI-6 for a more detailed analysis of changes in dynamics as a function of *J*.

Finally, the W2P function is also sensitive to the value of Δ, the half-width of the quenched noise distribution.

We provide in Supplementary Information (section J) an analysis of the response of the system to weak AC stimulation. Arnold tongues and plots of resonance frequency and amplitude of the system as a function of *J*, Δ and *a* show that the system is more sensitive for low values of *a* (slow synapses) and Δ, and for increased self-coupling.

### Stability properties

Although the fixed points of the extended model are the same as in the original MPR (they are not affected by the addition of the rate constants, as discussed above), the stability properties of fixed points are affected by the *a* parameter. For example, unlike with its traditional neural mass model analog, depending on the value of *a*, the NMM2 model exhibits oscillations (a Hopf Bifurcation) when the coupling constant is negative—an Interneuron Network Gamma (ING) interaction (see Figures SI-2 and SI-3). This has also been shown using first-order synaptic dynamics in (18). Furthermore, the excitatory network gamma version of the model (ENG) can also oscillate in the regime of low connectivity (small *J* > 0) and low excitability dispersion (small Δ, high synchrony)—see Figure SI-20.

### Resonance properties

We have carried out simulations of the response of the single population NMM2 model with excitatory self-coupling (*J* > 0) under the influence of a weak AC electric field perturbation, with *I*(*t*) = *P·E* = *A* sin(2*πft*). Simulations for *c* = *k* = 1 (which is equivalent working in the dimensionless unit system in Equation 99 in SI) are initiated by placing the system in its fixed point and then applying a weak perturbation,

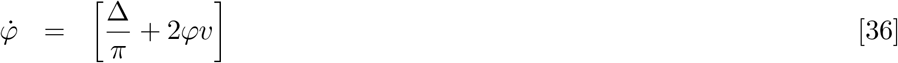

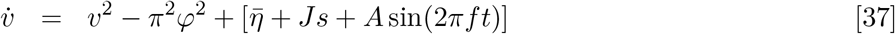

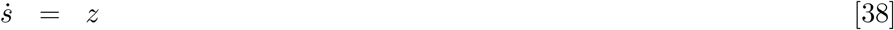

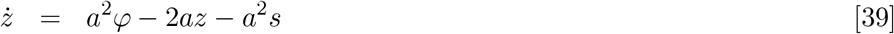

In this framework, *a* is the synaptic rate in membrane time units. Hence *a* ∼ 1 sets the system in a regime where the membrane and synaptic time scales are close to each other.

As we can see in Figure 5 and in the figures in supplementary material (section J), the system exhibits resonant peaks and *Arnold* tongues that shift to higher frequency, higher amplitude and smaller bandwidth as *J* increases for more details). The amplification factor at the resonant peak *σ/A* provides a normalized measure of the response of the system, measured as the standard deviation of potential divided by the input amplitude. The amplification factor weakens with increasing rate *a*, which corresponds to the system getting closer to the MPR model.

**Fig. 5.**
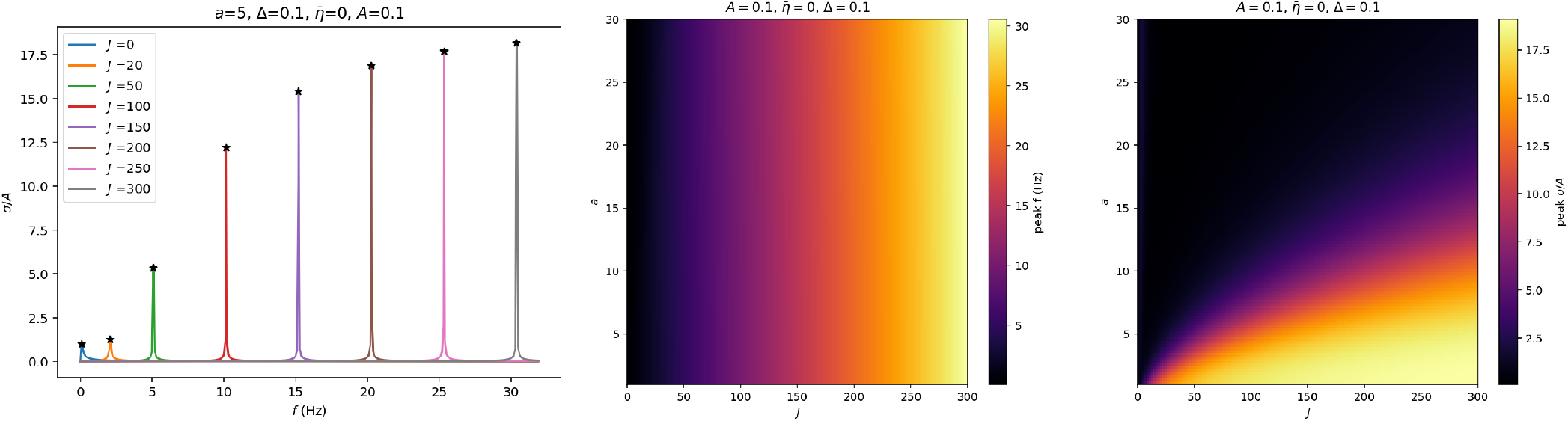
Sensitivity to input depends on *J* and rate constant in single population NMM2 model. Left: Normalized power as function frequency for different *J* values. Middle: Resonance frequency; Right: Bi-dimensional plot of resonant normalized peak power *σ/A* as a function of *J* and *a*.

This is seen in detail in Figure 5 and Figure SI-23, where resonance frequency *f*_*p*_ and amplitude (standard deviation *σ*) of the membrane potential are plotted as a function of *J*. Figure SI-24 displays further examples of the response amplitude as a function of *J* and Δ and as a function of *J* and *a*.

Finally, as the amplification factor increases, so does the transient time to reach steady state (both are related to the real part of the eigenvalues at the fixed point).

## 4. Discussion

The extended, unified model NMM2 brings together descriptions of synaptic transmission and neuron function based on mechanistic principles. This exact mean-field theory provides a physiologically robust description with parameters that can be related to physiological features. NMM2 features a supra-linear W2P function, exhibiting an increasing gain with growing inputs (and equivalently output firing rate). This means that effective coupling increases with increasing network activation, and as we saw, also with increasing self-coupling.

With regard to the self-coupling parameter *J*, note that in our formulation (following closely a biophysical interpretation) it has units of charge (see Equation 93). This reflects the fact that *J* encodes the effective connectivity of the average neuron to the rest of the network, i.e., the total number of connections/synapses the cell receives times the charge deposited by each during an action potential.

We have seen that in NMM2 the intrinsic population connectivity and other parameters play a key role in modulating the response of the system to its inputs. Translating this into the context of NMMs, this means that the lumped sigmoid parameters (*φ*_0_, *v*_0_ and *r*) should be adjusted to reflect the self-connectivity *J* and intrinsic noise half-width Δ of the cortical patch the mass model is representing. This is studied in more detailed, in SI Section K, where we show how to fit sigmoid parameters as a function of coupling for the case of slow synapses (the case most closely related to NMM). The most important change affects *φ*_0_, which increases monotonically with *J*.

Moreover, we can also see that there can be a delay in response to the input that depends on *J*—see Figure SI-4— or Δ (Figure SI-5). That is, in the examples shown, a loosely self-coupled population takes longer to react to the change input, and this delay is of a different nature than the postsynaptic delay: both the rate parameter *a* and *J*, Δ play a role in modulating the response-to-input delay. We can thus talk about a neural mass “inertia” that depends on these new parameters.

The analysis of the NMM2 framework also clarifies that the electrical coupling constant derived from neuron compartment models (26) should be directly used in NMM2 as is, without any scaling. The effective response of a single population to weak electric field perturbations will be modulated by the intrinsic population connectivity *J* or excitability (Δ, the quenched noise variance), as well as *a*, the rate constant. This suggests that if one wishes to work with the simpler NMM equations, some parameters, such as the sigmoid function parameters, may need to be adjusted, but we leave this analysis for further work.

The modulation of the effective coupling in NMM2 at the single population level provides a potential mechanism for the surprising sensitivity of networks to weak electric fields generated by brain stimulation or by neural tissue itself (ephaptic interaction (27)). It also highlights the fact that in the more general setting of multi-population dynamics, connectivity within and across populations play an important role in modulating the sensitivity to inputs. This may also explain why stronger electrics fields are needed to induce measurable effects in in-vitro (low, disrupted connectivity) or small animal studies (smaller cortical patch areas) as compared with humans.

A tightly coupled population in the right regime can display more sensitivity to weak external inputs (see the SI section J for examples), although, in general, this will depend on the state of the network and factors such as the type of E-I balance. With regard to this, current evidence favors *loose balance* in the sensory cortex (defined to be when net input remaining after cancellation of excitation and inhibition is comparable in size with the factors that cancel), for example, but in the motor or frontal cortex the case is less clear, with *tight balance* also having been proposed (when the net input is very small relative to the canceling factors) (28). Presumably, the weak perturbation effects of tES would be more important in the tight balance regime.

In transcranial electric stimulation (tES), the electric field generated on the cortex is of the order of 1 V/m (29), which is known to produce a sub-mV membrane perturbation (25, 30). Yet, it is of mesoscopic nature, with a spatial scale of several centimeters. The main characteristic of the exogenous macroscopic electric field generated by tES is its small magnitude, low temporal frequency, moderate spatial correlation scales (> 1 cm), a long application time (typically twenty minutes to one hour) and application with repeated sessions when long-term plastic effects are desired. Endogenous fields are similarly characterized by weak field magnitudes over mesoscopic scales (27). As we have seen in the analysis of an exact mean-field theory, weak but spatially uniform fields can affect the dynamics of densely connected neuron populations exposed.

The NMM2 formalism introduces new dynamical features, even in simple models. The MPR model, which can be seen as an NMM2 in the limit of very high rate constant (fast synapses), already displays chaos (15). We have seen here that NMM2 can produce rich dynamics, with features such as oscillations in a single population with constant input, as well as bursting in a simple two population model (SI sections C, D, and E). We leave the detailed analysis of these and other systems in the NMM2 framework for further work.

## 5. Conclusions

Based on a recently derived exact mean-field theory and decades of work on neural mass modeling, we provided an extension of first that brings together the features in each with a solid mechanistic basis. The resulting model is limited, as it is based on a simple neuron population model (QIF with uniform self-connectivity), but is an exact mean-field theory endowing with unambiguous meaning each of its parameters and variables. The analysis of the resulting system and comparison with classical NMMs sheds light on network phenomena mediated by self-connectivity and the physiological basis of semi-empirical parameters in NMMs.

## ACKNOWLEDGMENTS

This work has received funding from the European Research Council (ERC) under the European Union’s Horizon 2020 research and innovation programme (grant agreement No 855109) and from FET under the European Union’s Horizon 2020 research and innovation programme (grant agreement No 101017716). This work has benefitted from multiple discussions with my collaborators, Paul Clusella, Elif Koksal and Jordi Garcia-Ojalvo.

## Supplementary Information

### A. Dipole nature of electric field interaction in compartment model of a neuron in the linear regime

Here we show that in the linear regime (far from threshold) or when the electric field is weak, the effect of an electric field on a neuron compartment, or its average over a set of compartments, can be written in the form *δV* = *λ·E*, where *λ* is a vector. This is the so-called “lambda-E” model used in transcranial electrical stimulation models (29, 31).

The effect of an external field can be represented in a neuron compartment model by an axial current that results from the potential difference induced by the field along the fiber associated with a compartment (see, e.g., (26) and references therein),

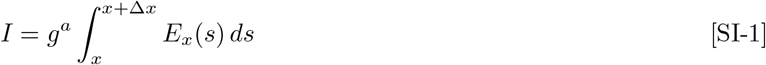

where *E*_*x*_ is the field along the fiber. We can rewrite this as *I* = *g*^*a*^∫_*f*_ **E** · *d***l** with the line integral along the fiber compartment. If the field is constant along the fiber we can express this simply as *I* = **E · u**. The vector **u** is a vector parallel to the line that ends in the compartment of interest and originates in the connected one(s), and thus pointing into the compartment (see Figure SI-1). If the E field is aligned with this direction, we get a positive current into the compartment. If the compartment of interest is *k* and the connected compartment is *j*, we refer to the vector **u**_*k,j*_ parallel to **x**_*k*_ − **x**_*j*_ (the coordinates of the compartments). The superscripts *r* and *a* refer to radial and axial components and conductivities.

**Fig. SI-1.**
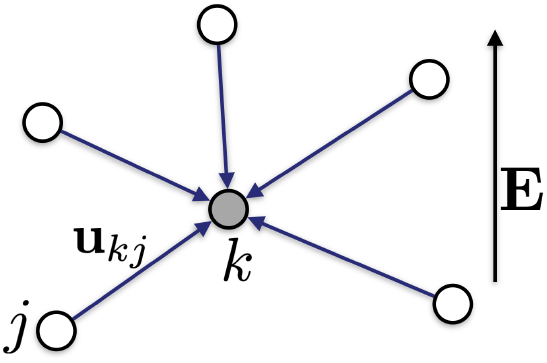
Compartments and vectors for cable equation in E field. Here we represent a fiber by a node and and a set of directed edges.

From conservation of charge (the sum over *j* is over connected compartments, and *i* over membrane currents), and assuming that the field is uniform and thus independent of the compartment *k*, we obtain

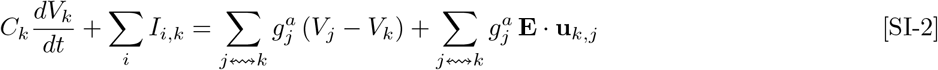

The sum Σ_*j*↭*k*_*Q*_*j*_ ≡ Σ_*j*_*c*_*kj*_ *Q*_*j*_ denotes the sum over compartments connected to the *k*th one, and can be expressed via the compartment connectivity matrix *c*_*kj*_, with value of 1 if compartment *j* is connected to *k*, zero otherwise. The sum on the left hand side is over ionic membrane currents out of the compartment.

We can simplify this and also pull the constant electric field out the sum,

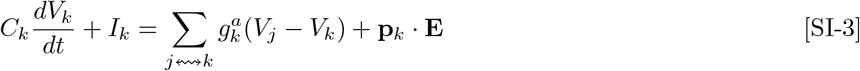

where *I*_*k*_ = Σ_*i*_*I*_*i,k*_, the *k* sum is over connected components to the *k*th compartment and where

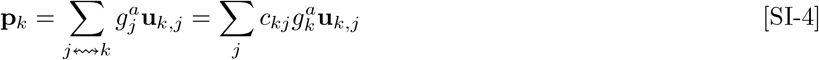

The equation for a single compartment in steady state is thus of the form

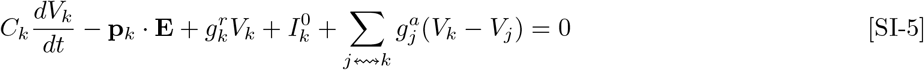

where **p**_*k*_ is the dipole vector corresponding to the *k*th compartment, and 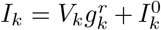, with *I*^0^ the sum of currents associated to the reversal potentials, and 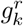 the axial (transmembrane) conductivity in the linear regime.

Thus, we see that the behavior of each compartment is characterized by its own dipole and the interaction with other compartments.

Equation SI-5 is linear and can be expressed in matrix form, with the compartment array notation *V* = (*V*_*k*_) = (*V*_1_, *V*_2_, …, *V*_*N*_), *C* = diag(*C*_*k*_) = diag(*C*_1_, *C*_2_, …, *C*_*N*_), and (**p**_*k*_ · **E**) = (**p**_1_ · **E**, …, **p**_*N*_ · **E**),

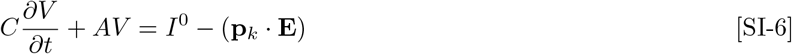

and with the matrix *A* given by

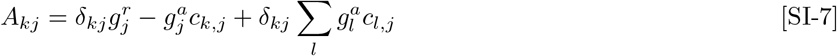

In steady state 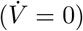, the solution to this equation is

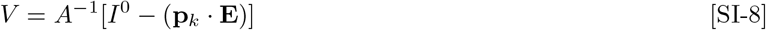

or

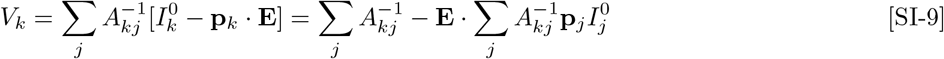

where the second equality is a result of linearity of the *A* matrix, i.e., the fact that we are dealing with a linear equation. Finally, we can write the solution as

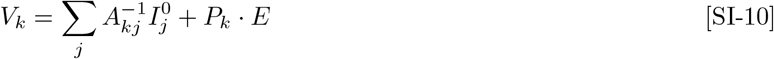

This equation shows that the response of the compartment to an electric field is of dipole form, with the total dipole *P*_*k*_ resulting from a superposition of dipoles,

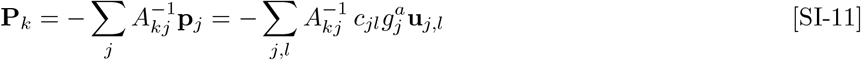

Furthermore, we observe that any linear function of compartment potentials will also display, by further superposition, a dipole response. This includes, for example, the average membrane perturbation for apical dentrite compartments, or soma.

What happens in the **nonlinear case**, when the neuron is not at baseline? We can write (steady state)

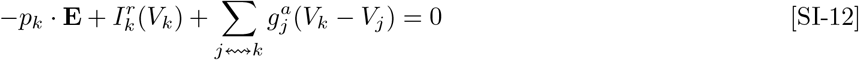

with 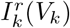 a nonlinear function, or, in a fashion analogous to the discussion above, in matrix form

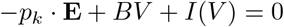

or

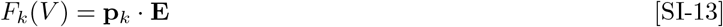

for a nonlinear operator *F*. This means we cannot pull out the E field out of the inverse. However, if the electric field is very weak, we can carry out the analysis above with the linearized version of Equation SI-13 by expanding it around the potential with zero field, with the same conclusion.

### B. Stability analysis for single population NMM2 model

We analyze here the fixed points and their stability of the NMM2 single population model, showing it displays a Hopf bifurcation for some parameters choices.

We start from the equations

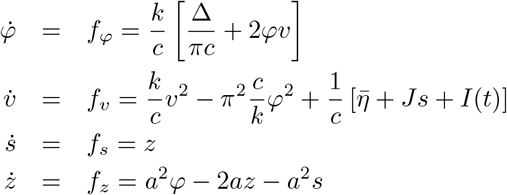

Fixed points, as described in the methods section, Equation 33, are determined by

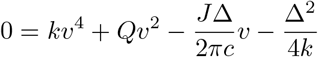

We can express *Q* as a function of the other parameters,

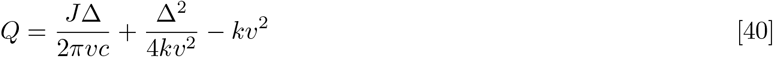

The Jacobian is

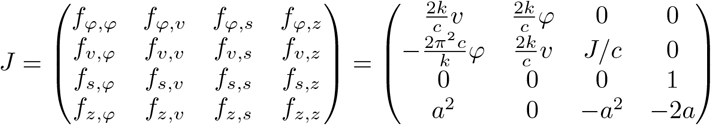

To assess stability of a fixed point we compute the eigenvalues of the Jacobian at that point, and if any points have a positive real component they are marked as unstable.

**Fig. SI-2.**
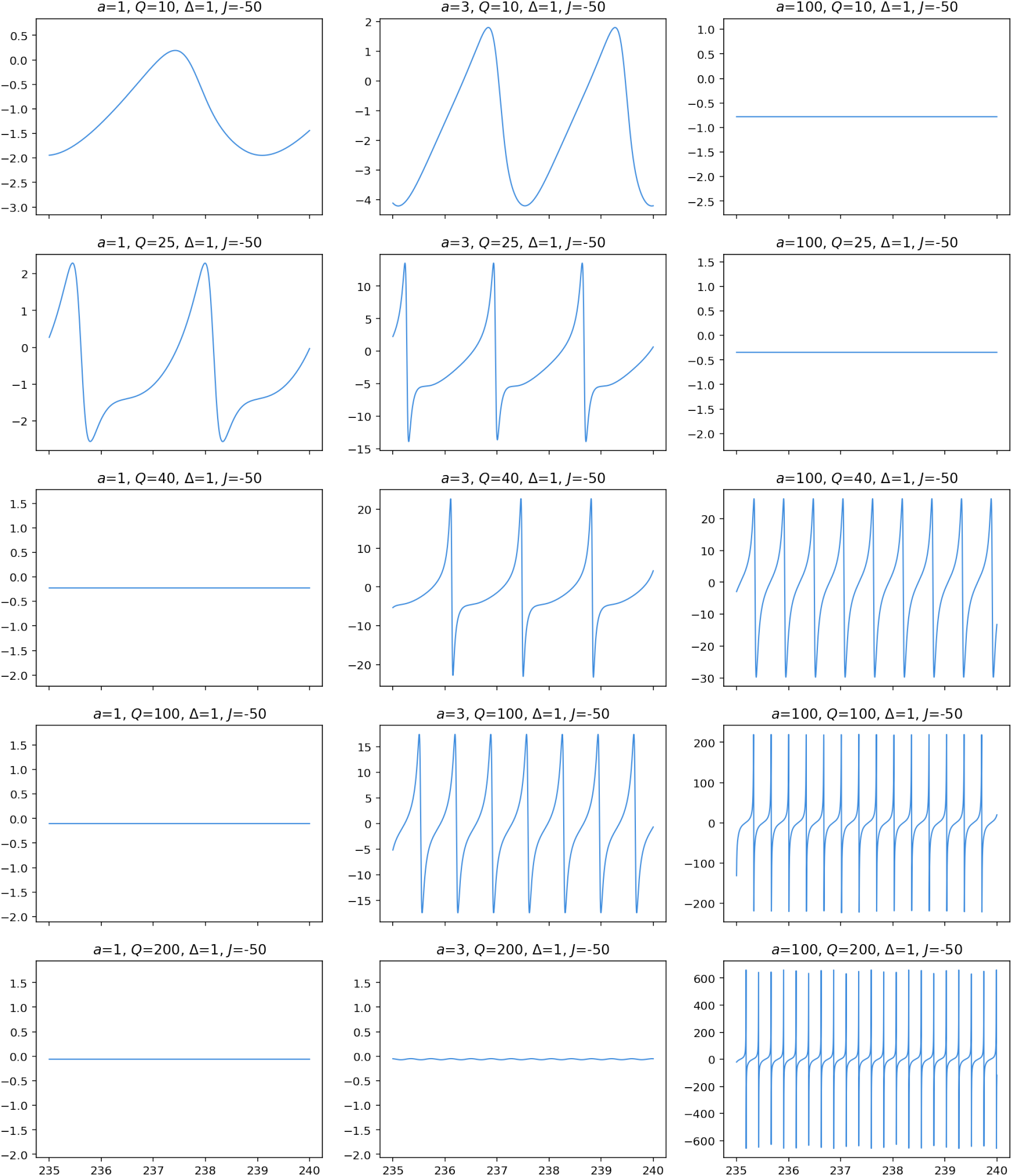
Sample dynamics for self-inhibitory single population system with *J* = − 50 and Δ = 1. Graph axis are time (horizontal) and mean membrane potential (vertical).

**Fig. SI-3.**
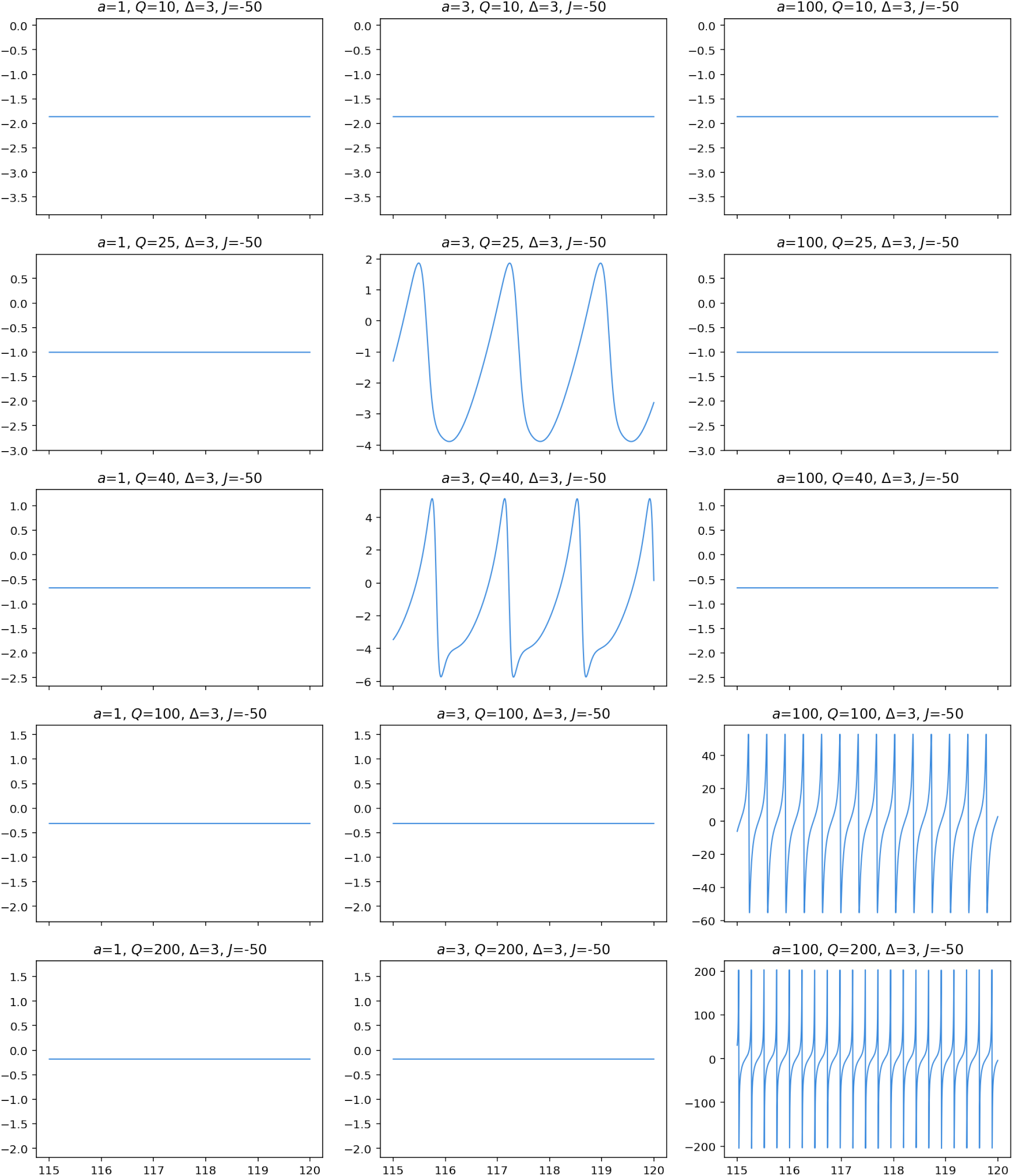
Sample dynamics for self-inhibitory single population system with *J* = − 50 and Δ = 5. Graph axis are time (horizontal) and mean membrane potential (vertical).

**Fig. SI-4.**
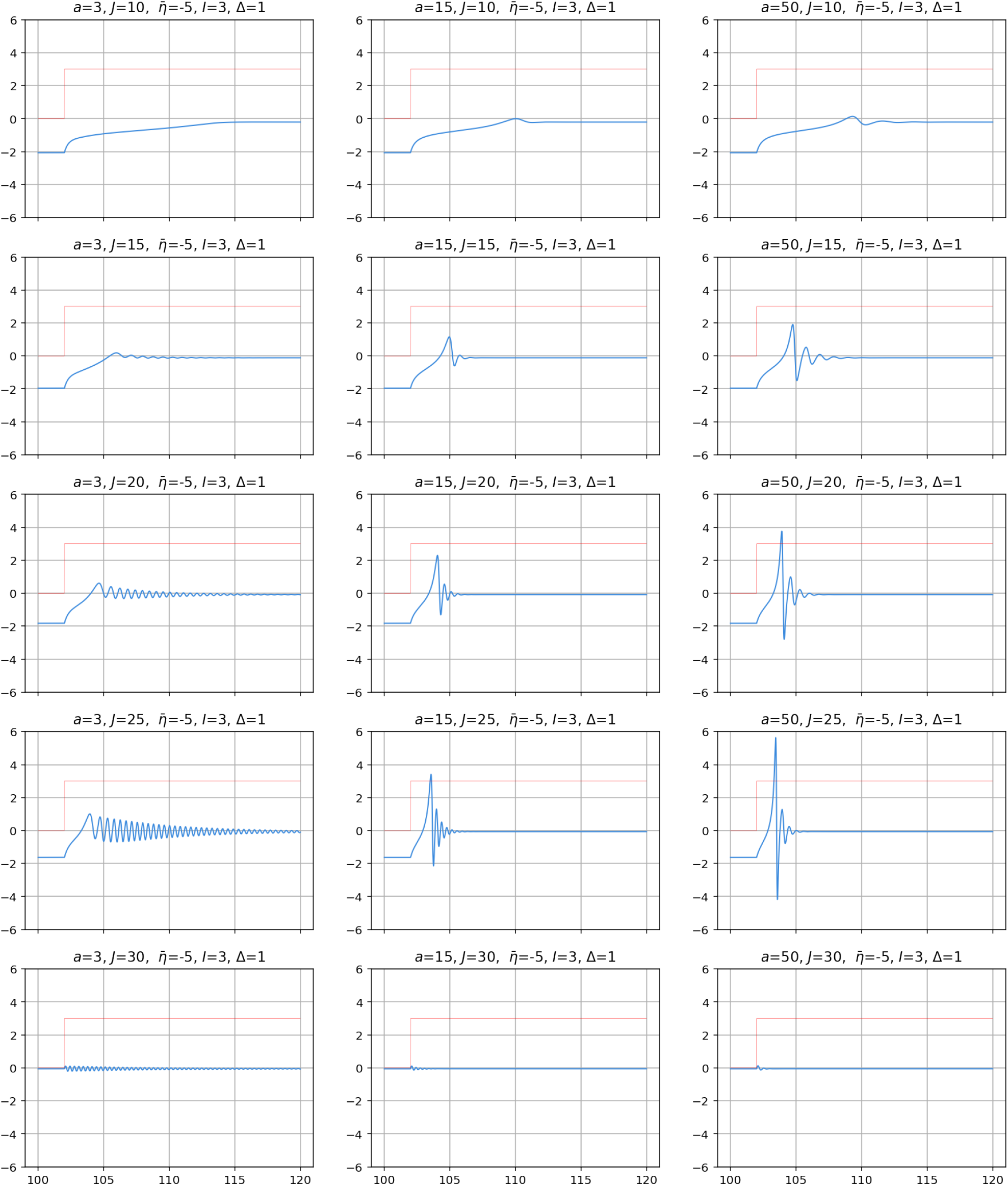
Response of excitatory (*J* > 0) single population to step input function (red) as a function of self-coupling. **Here the total input is** 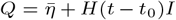. Note how the delay in response of the population to the input (neural “mass inertia”) varies with both *J* and *a*. Graph axis are time (horizontal) and mean membrane potential (vertical).

**Fig. SI-5.**
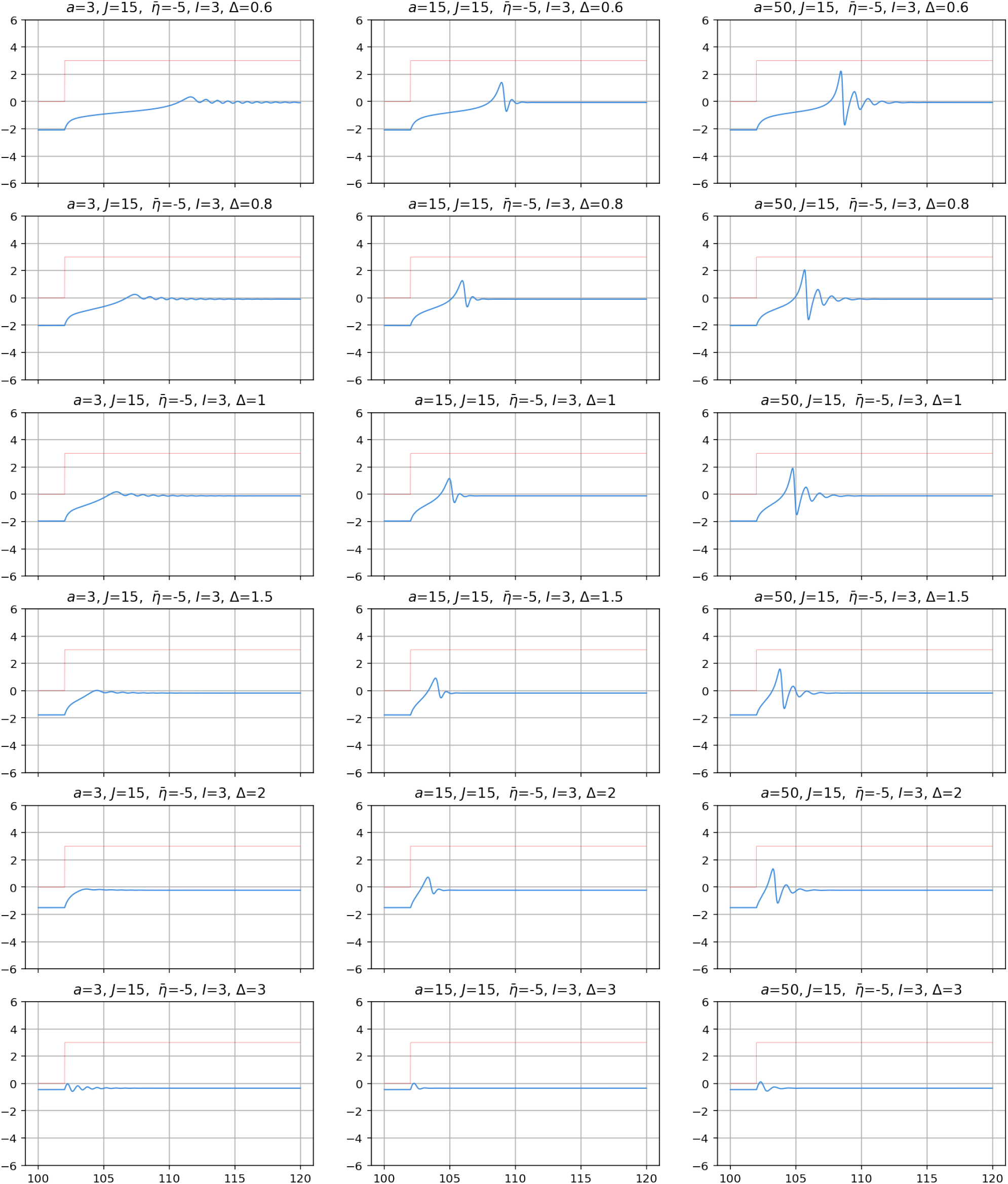
Response of excitatory (*J* > 0) single population to step input function (red) as a function of Δ. **Here the total input is** 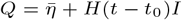. Note how the delay in response of the population to the input (neural “mass inertia”) varies with both Δ and *a*. Graph axis are time (horizontal) and mean membrane potential (vertical).

**Fig. SI-6.**
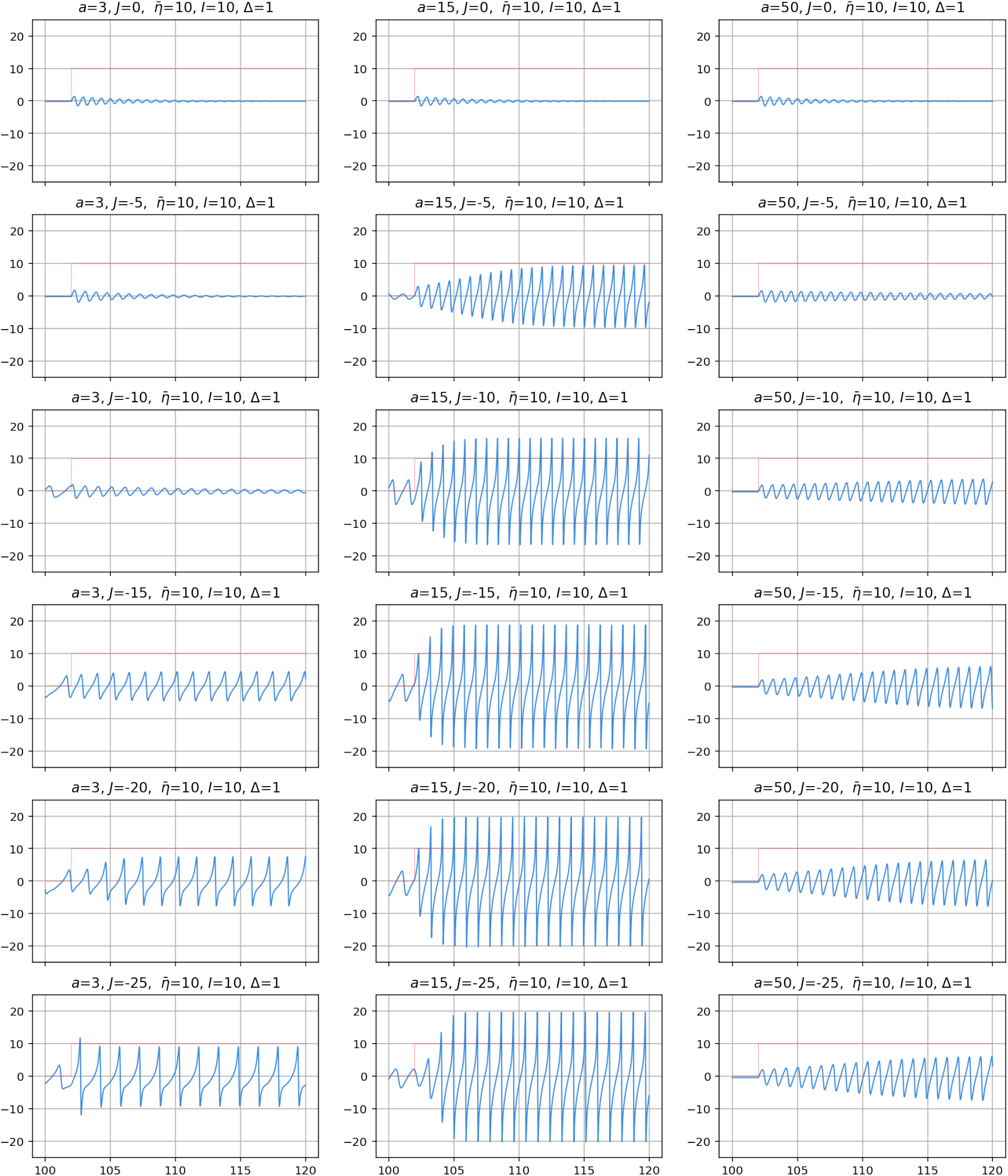
Response of inhibitory (*J* < 0) single population to step input function (red) as a function of self-coupling. Here the total input is 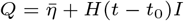. Graph axis are time (horizontal) and mean membrane potential (vertical).

### C. Sample dynamics for simplified NMM2 E-I model

#### Two population excitation-inhibition (E-I) model

Excitation-inhibition models such as pyramidal-interneuronal network gamma (PING) (23, 32–34) are often used to produce fast oscillations through a single Hopf bifurcation and are the simplest circuits in NMM capable of spontaneously oscillating under a constant input (35). They consist of an excitatory and an inhibitory population coupled together that can represent, for example, pyramidal and fast inhibitory cells in the cortex. We provide here an analog circuit in the NMM2 formalism. For the purposes of analysis, we simplify this system to have a simple constant input *Q*, and common parameters *a*, Δ, and we neglect self-coupling terms (Γ_*nn*_ = 0, see Figure 1). The equations for this system are

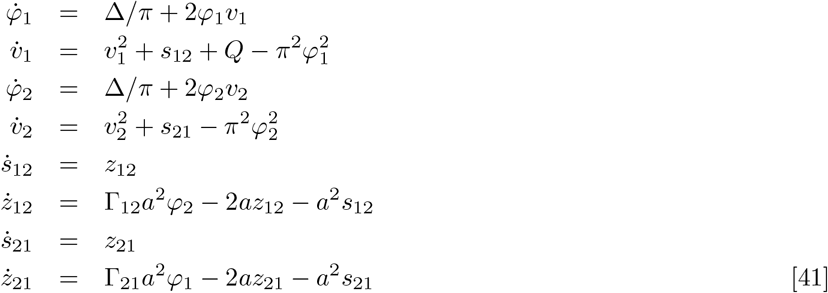

This system of equations exhibits rich dynamics. Figure SI-7 provides some sample dynamics, which now include bursting. Figure SI-12 displays simple bifurcation diagrams (starting from the same initial conditions (0,-2,0,0) and eliminating a transient of 10 seconds) to study the effect of changing the rate parameter *a*. The tails of these diagrams are probably noise from long transients. Figure SI-13 displays the increased sensitivity of the model to input when the coupling is increased while maintaining the E/I balance (here the ratio of the Γ_21_*/*Γ_12_ constants).

**Fig. SI-7.**
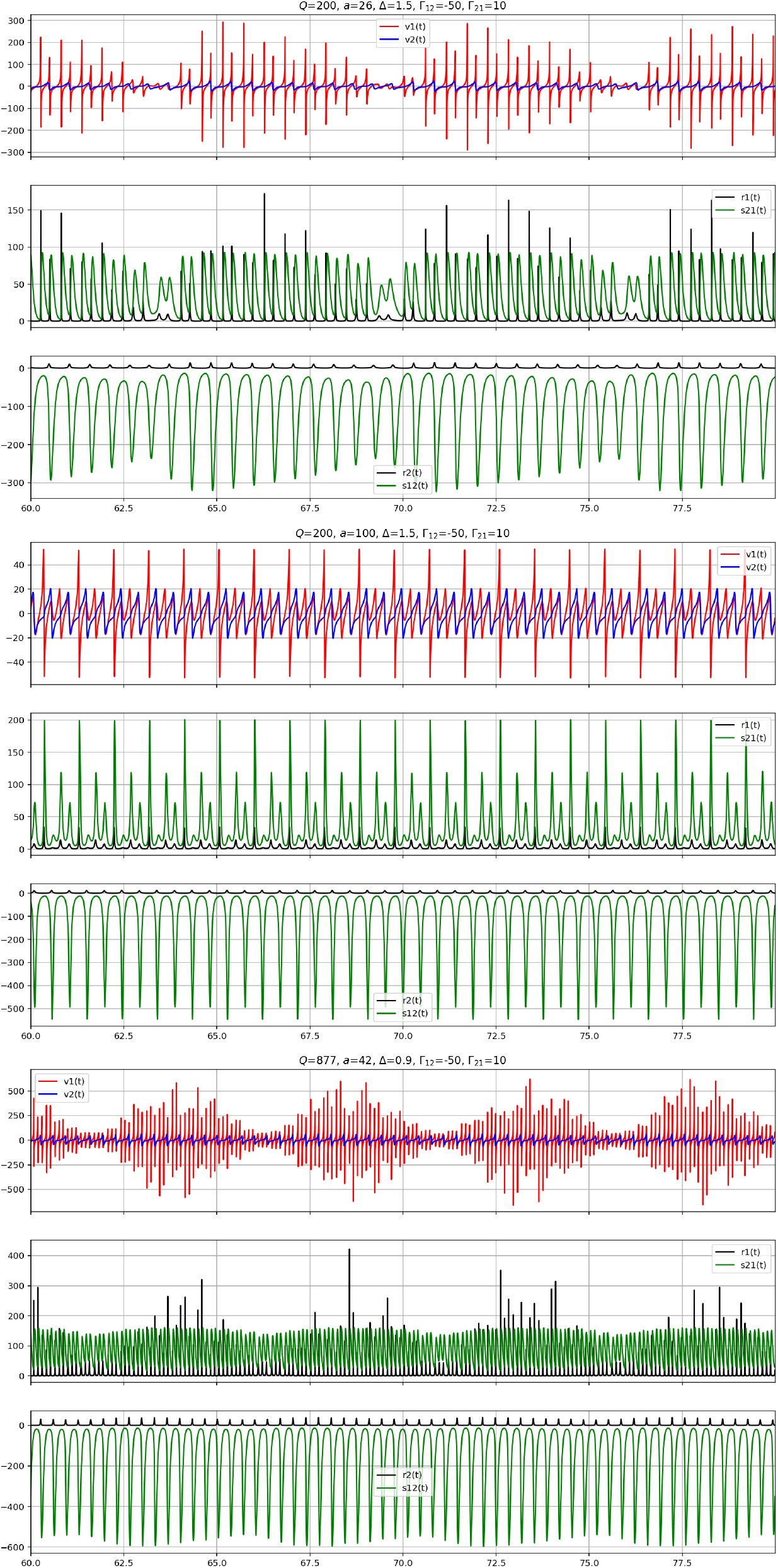
Sample dynamics for the NMM2 two population E-I model. Horizontal axis is time (horizontal).

**Fig. SI-8.**
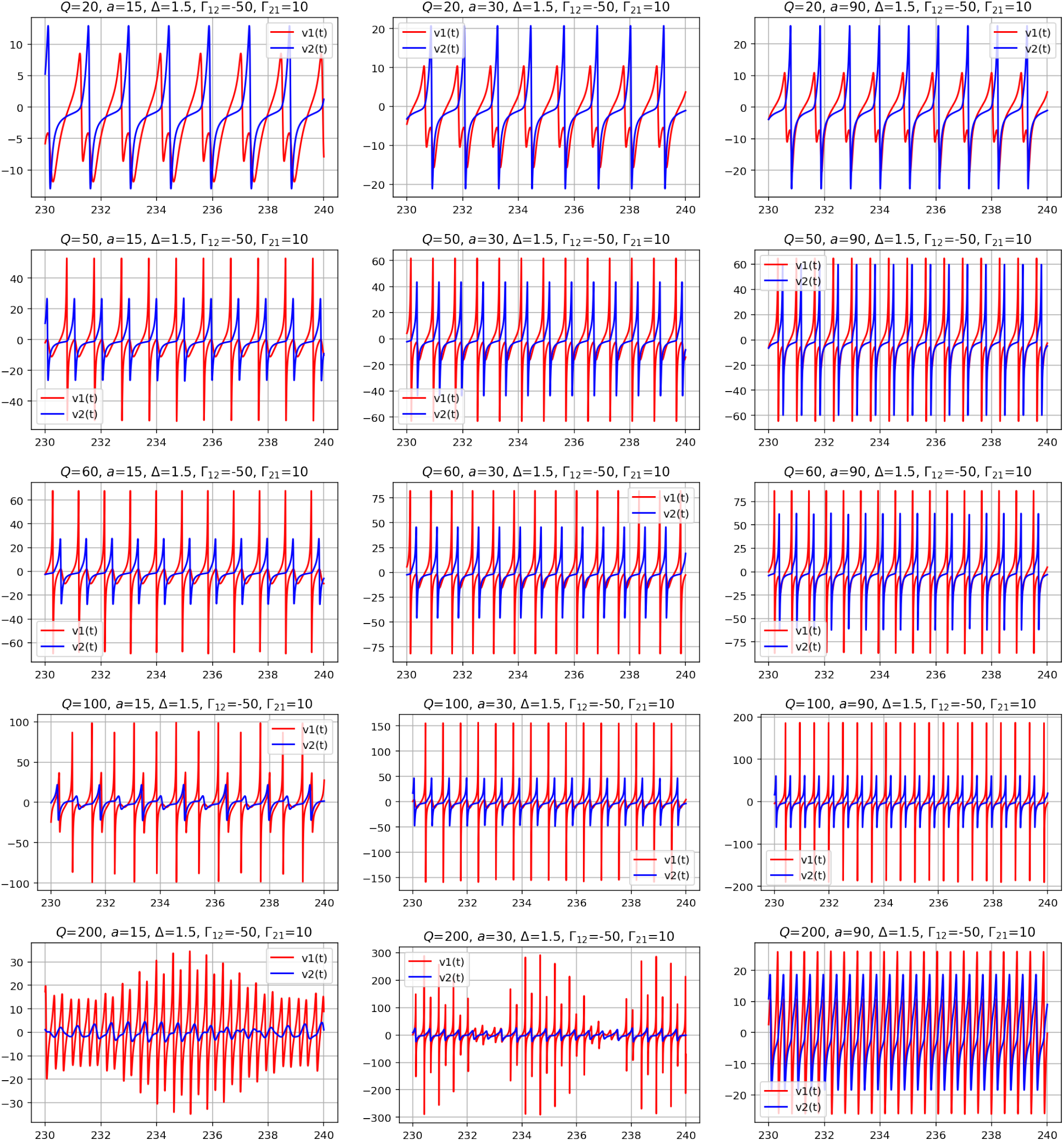
Sample dynamics for the NMM2 two population E-I model. Graph axis are time (horizontal) and mean membrane potential (vertical).

**Fig. SI-9.**
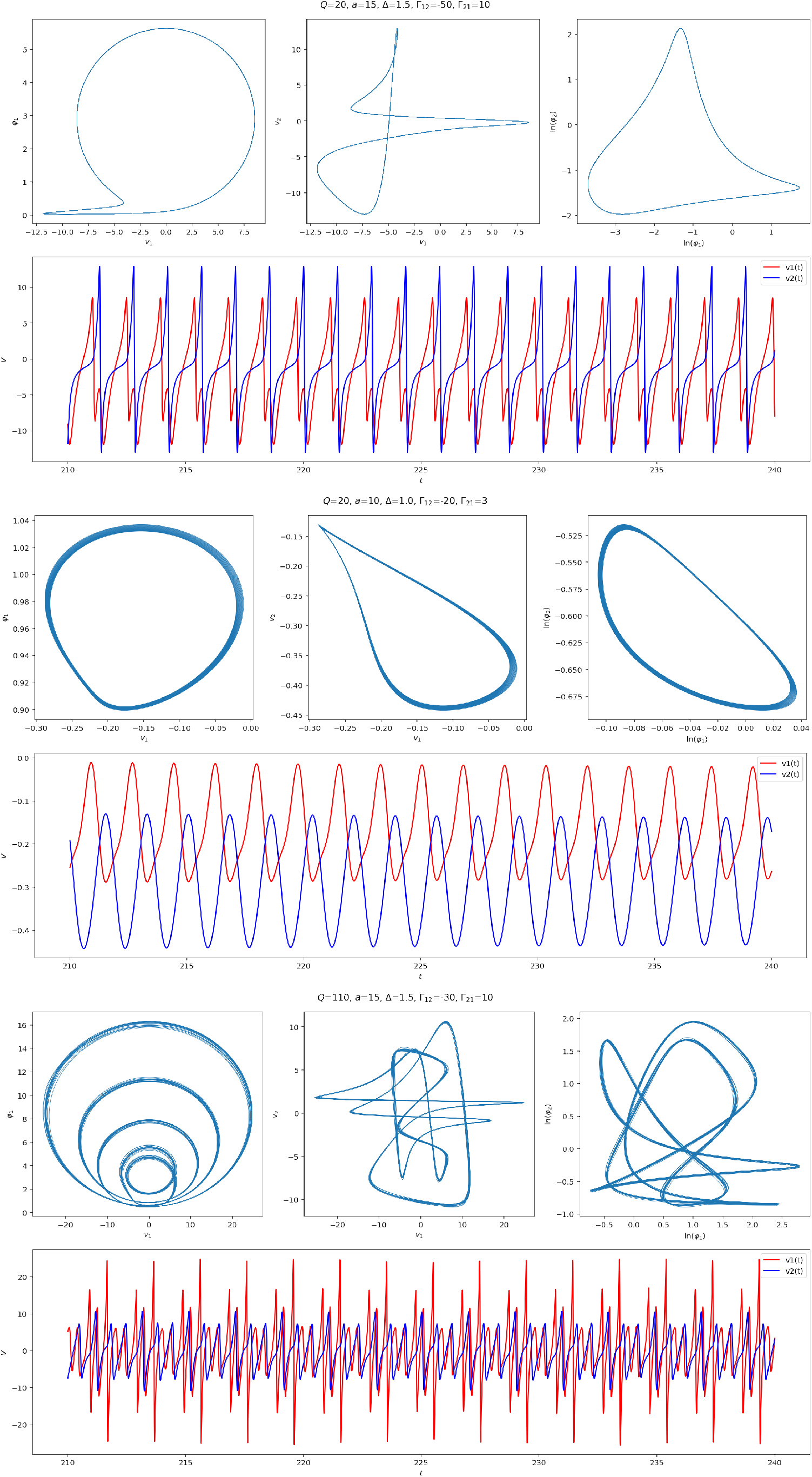
Interesting dynamics for the NMM2 two population E-I model (I).

**Fig. SI-10.**
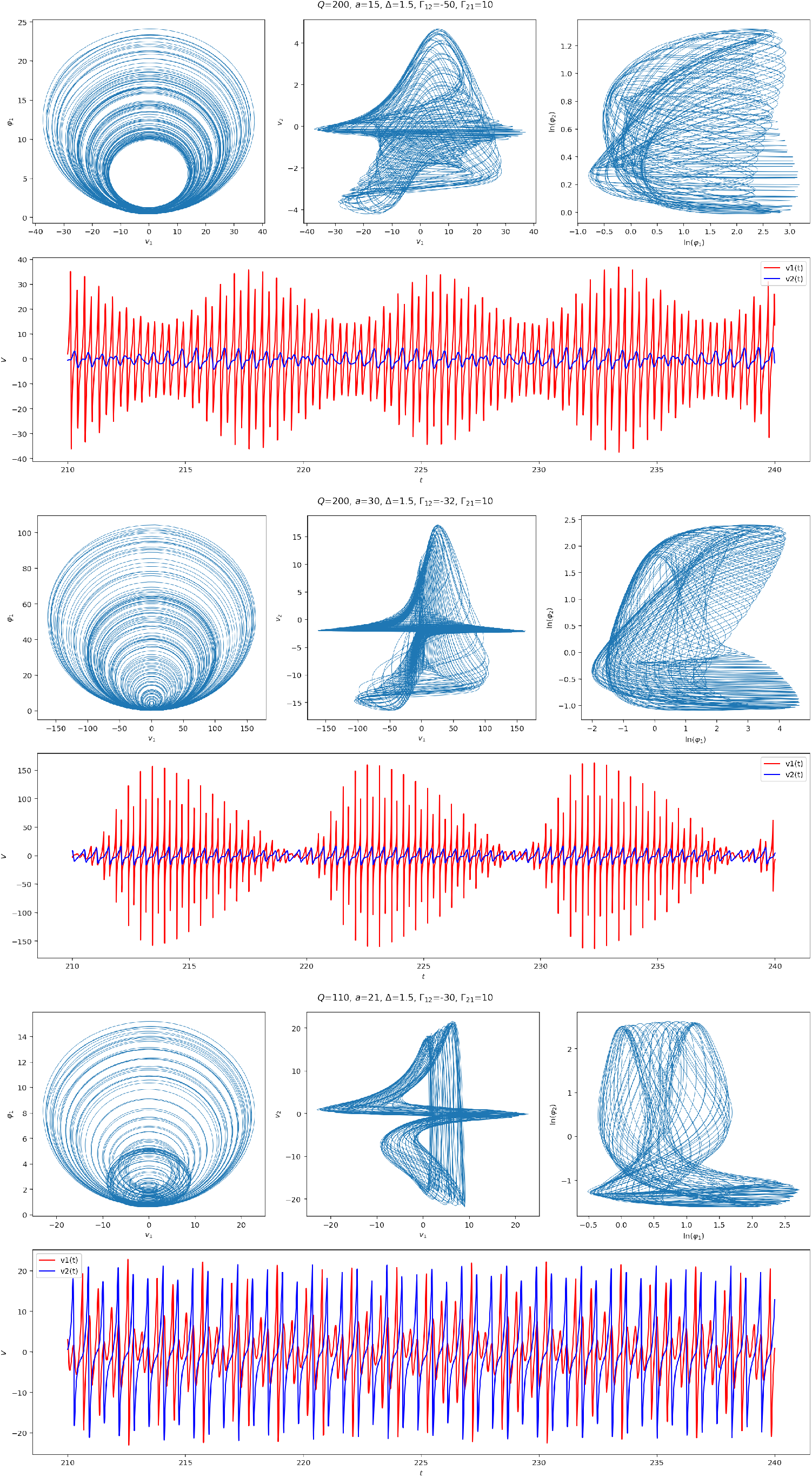
Interesting dynamics for the NMM2 two population E-I model (II).

### D. Bifurcation diagrams for simplified NMM2 E-I model

Here we provide some bifurcation diagrams for the simplified E-I model, by simply plotting the dynamics for each step of the bifurcation parameter (input current).

**Fig. SI-11.**
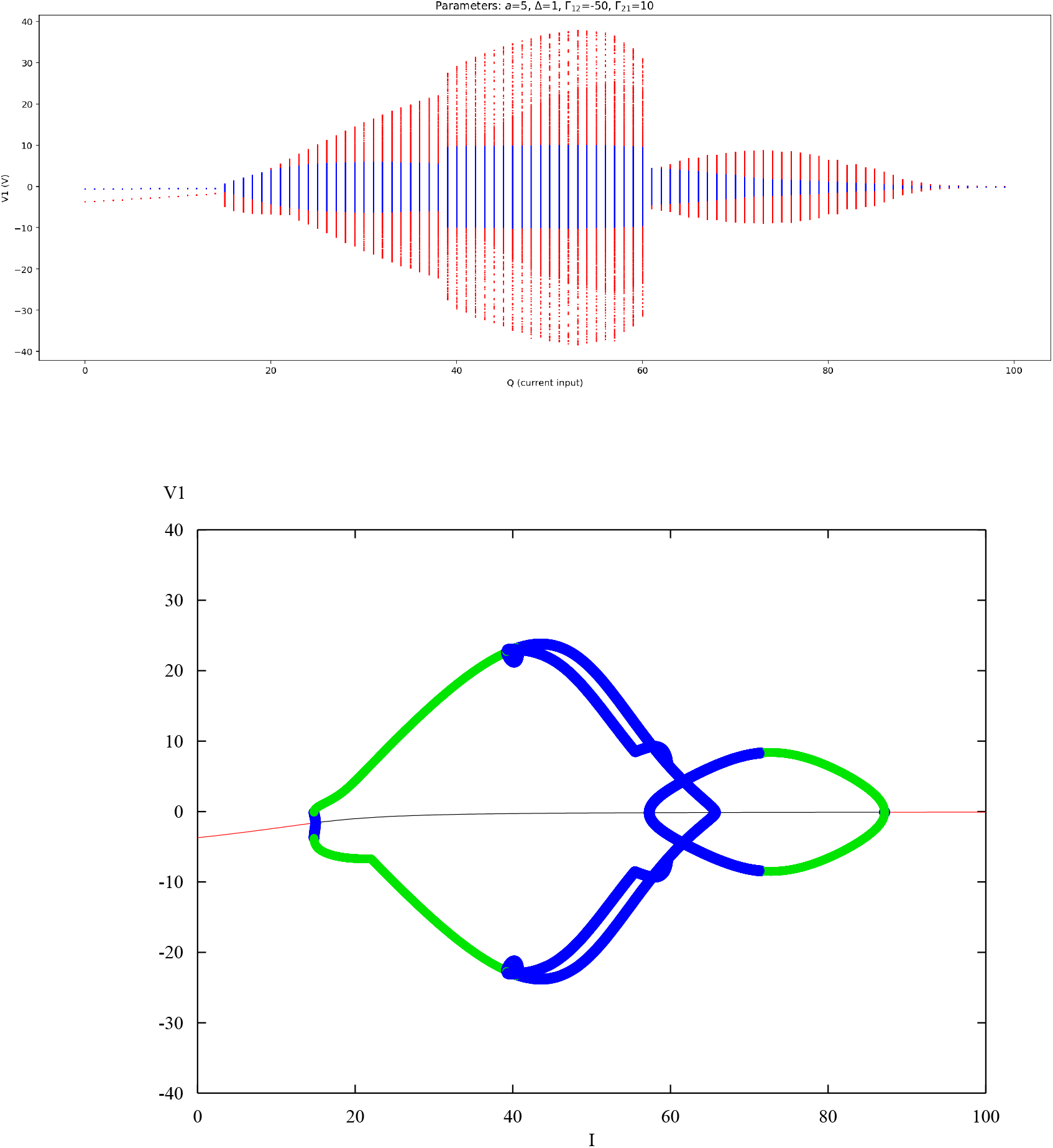
Bifurcation analysis of NMM2 E-I model for *a* = 5 with XPP-AUTO (bottom, see 07.5 MPR bifurcation.ipynb). The diagram exhibits multiple bifurcation points, including Hopf and period doubling points.

**Fig. SI-12.**
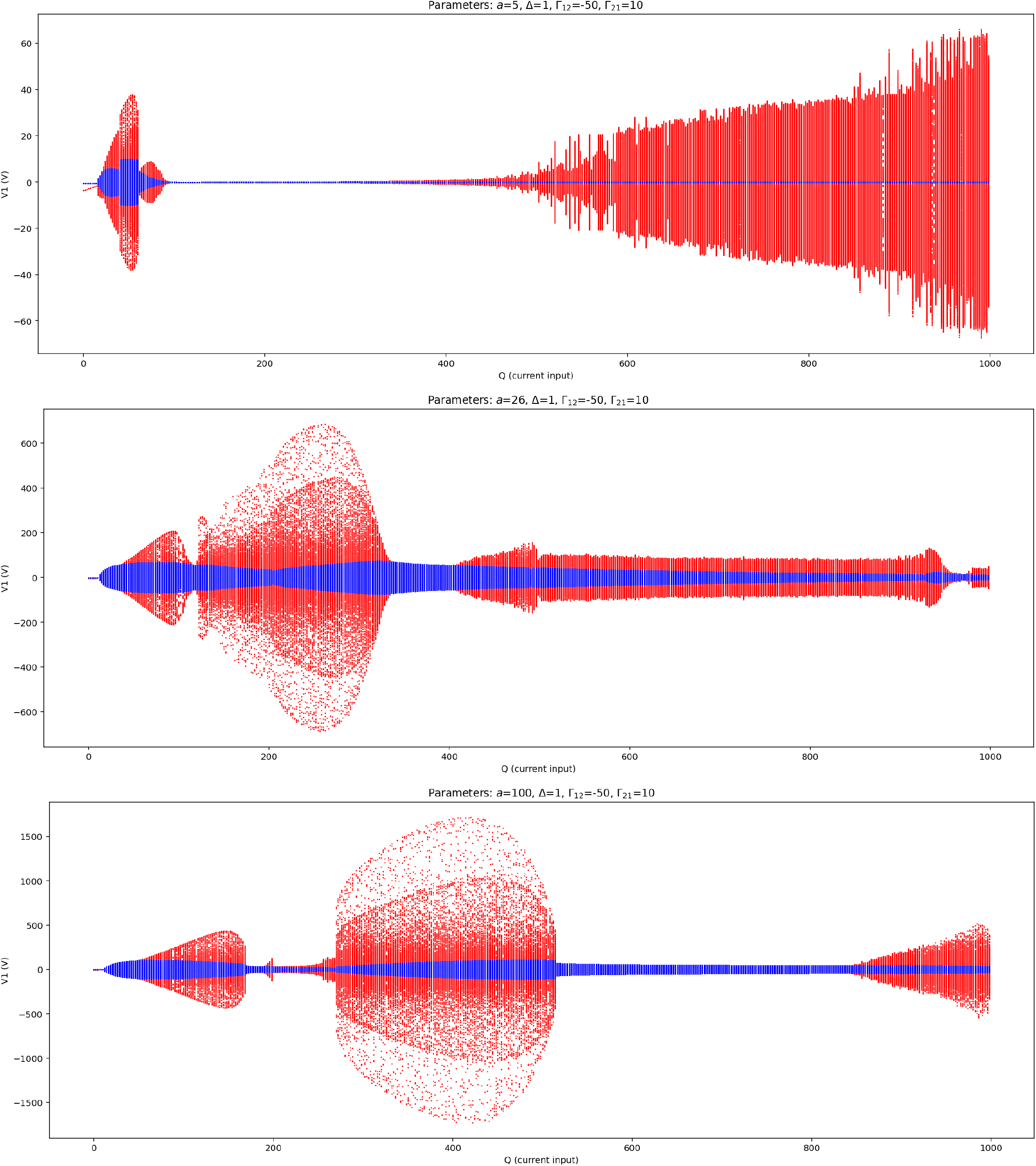
Bifurcation analysis of NMM2 E-I model for different values of *a* (5,25,100).

**Fig. SI-13.**
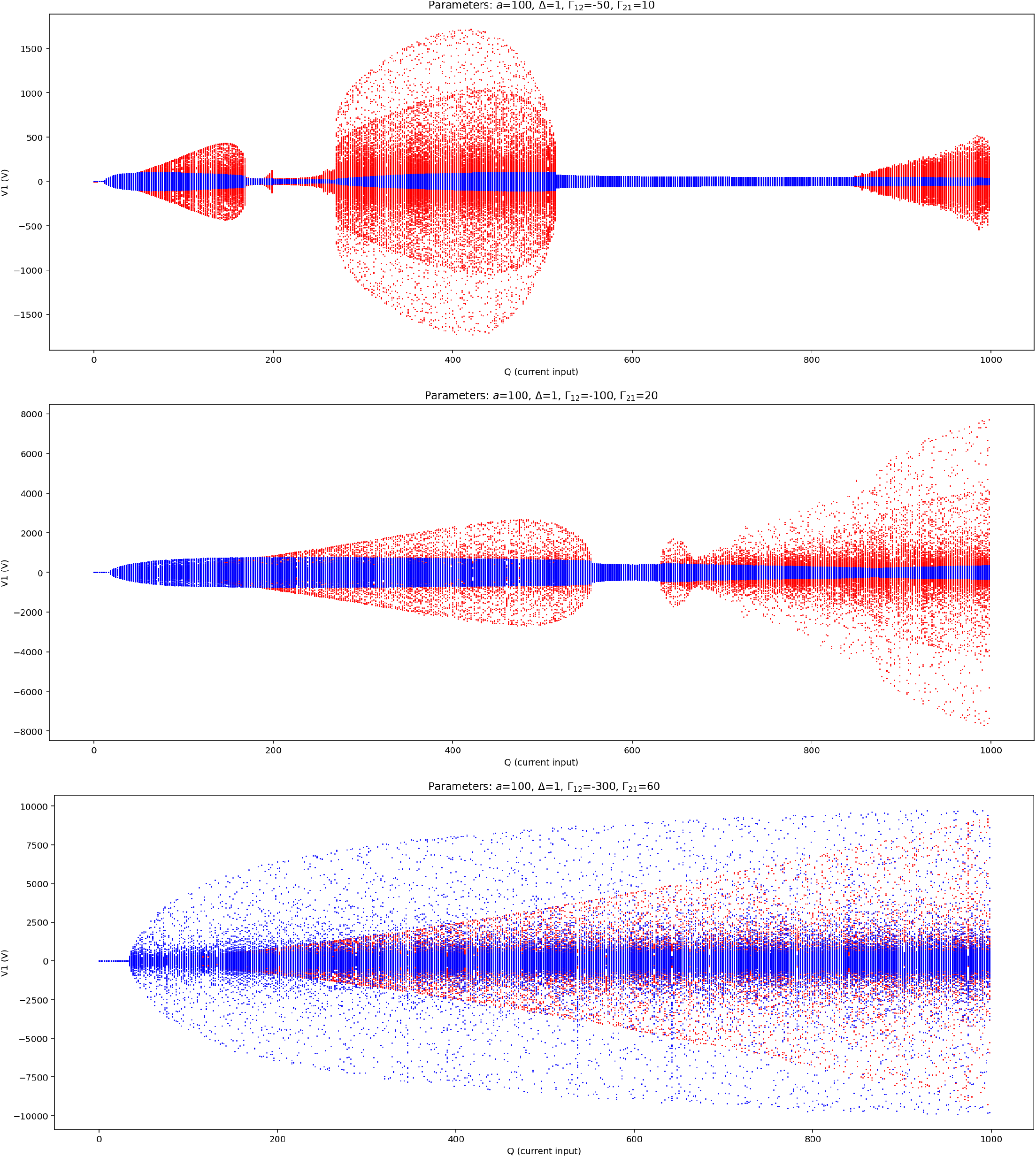
Bifurcation analysis of NMM2 E-I model for different values of coupling gain (Γ increasing from top to bottom, with same E/I ratio). Note the increase in voltage as the gain scaling is increased.

### E. More general NMM2 E-I model

Here we retain more features of the full two population model. The system of equations becomes — first the firing rate equations,

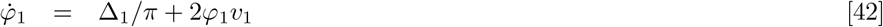

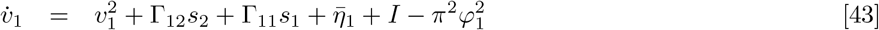

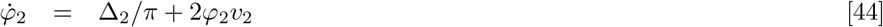

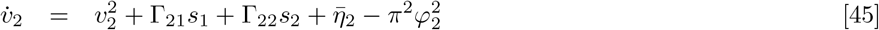

and the synapse equations, assuming the type of synapse is fixed for each source neuron,

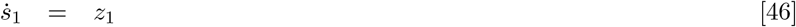

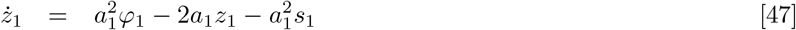

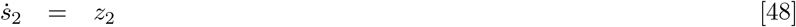

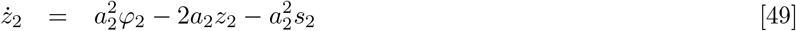

**Fig. SI-14.**
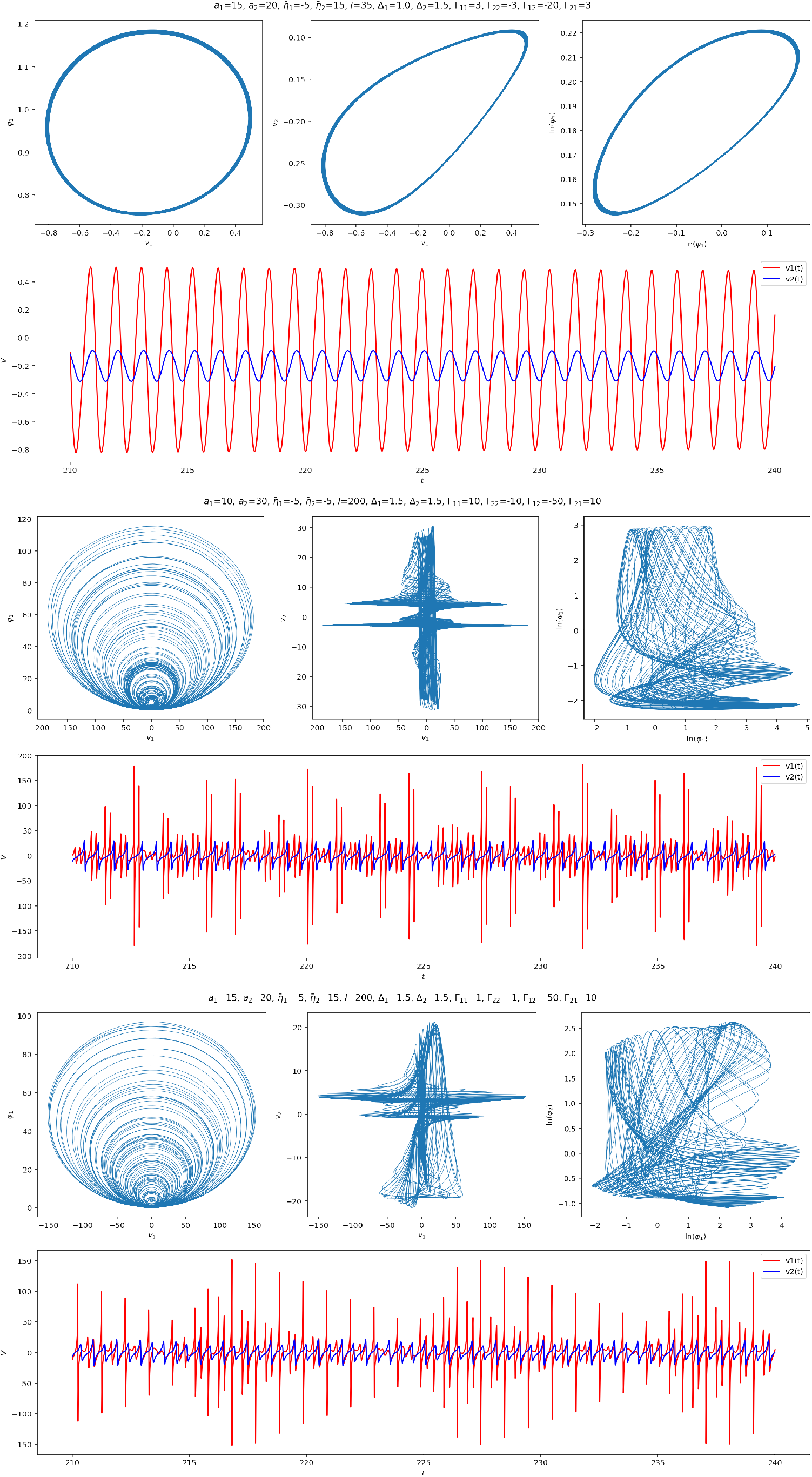
Interesting dynamics for the NMM2 more general two population E-I model (III).

### F. Realistic parameters for QIF model

Our starting point is the QIF equation,

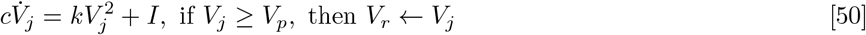

The units adapted to the scale of the problem are provided in Table 1. All parameters are typical from the literature, except *k*, which is adapted to produce reasonable firing rates. The resulting response function is provided in Figure SI-15.

**Fig. SI-15.**
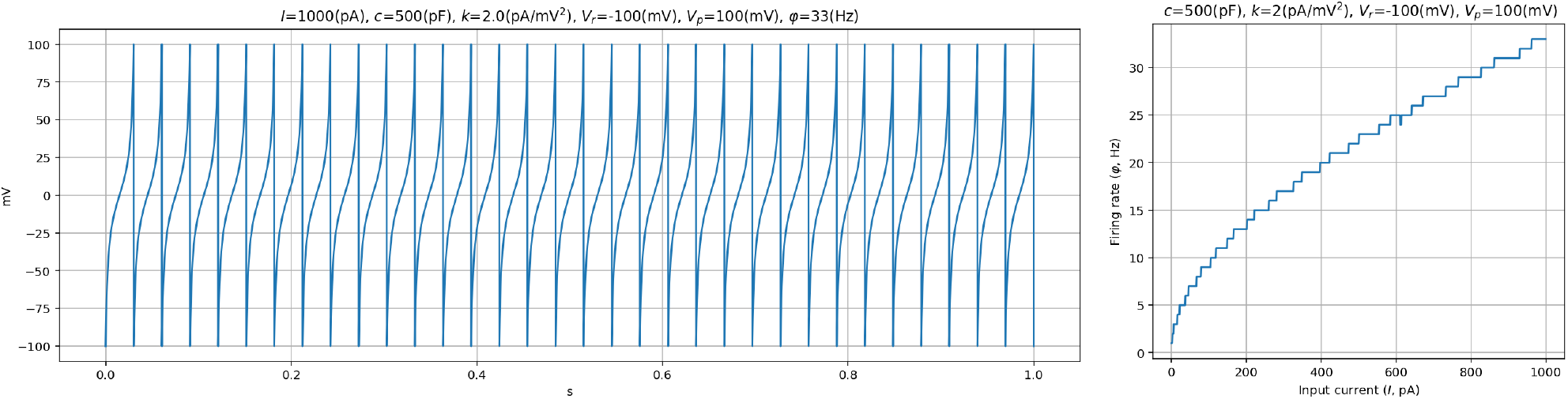
Sample dynamics and response function for selected QIF parameters (see (36) for comparison).

**Table 1.** **QIF variables, parameters with typical values and units (see, e**.**g**., **(36). The value for** *k* **has been determined by finding reasonable behavior in the QIF model (firing rate)**.

| Symbol | Units | Typical values |
| --- | --- | --- |
| $V$ | mV | 0–100 mV |
| $V_r$ | mV | -100–50 mV |
| $V_p$ | mV | 50–100 mV |
| $I$ | pA | 0–1000 nA |
| $c$ | nF | 200–1000 pF |
| $k$ | pA/mV<sup>2</sup> | 1–5 pA/mV<sup>2</sup> |

### G. Realistic parameters for NMM2 model

The values related to the underlying QIF model are provided in the previous section.

The full equations are

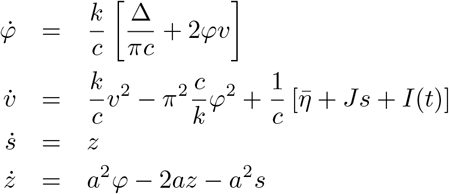

and the extra parameters are 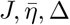 (which replace sigmoid parameters) and the synaptic rate *a*.

With regard to *J*, this is the average charge delivered by a an action potential (the integral of the PSC). From (37), we can estimate *J* to be about 50 ms · 500 pA, or about 25 pC.

The parameters, variables and NMM2 units are provided in Table 2. The resulting response function is provided in Figure SI-15.

**Table 2.** NMM2 variables, parameters with typical values and units — see, e.g., (36, 37).

| Symbol | Units | Typical values |
| --- | --- | --- |
| $V$ | mV | 0–100 mV |
| $V_r$ | mV | -100–50 mV |
| $V_p$ | mV | 50–100 mV |
| $I$ | pA | 0–1000 nA |
| $c$ | nF | 0.2–1.0 nF |
| $k$ | pA/mV<sup>2</sup> | 1–5 pA/mV<sup>2</sup> |
| $v$ | mV | -50–50 mV |
| $\varphi, s$ | Hz | 0–100 Hz |
| $J$ | pC | 5–50 pC (37) |
| $\bar{\eta}$ | pA | -500–500 pA |
| $\Delta$ | pA | (0,500] pA |
| $a$ | Hz | 20–500 Hz |

**Fig. SI-16.**
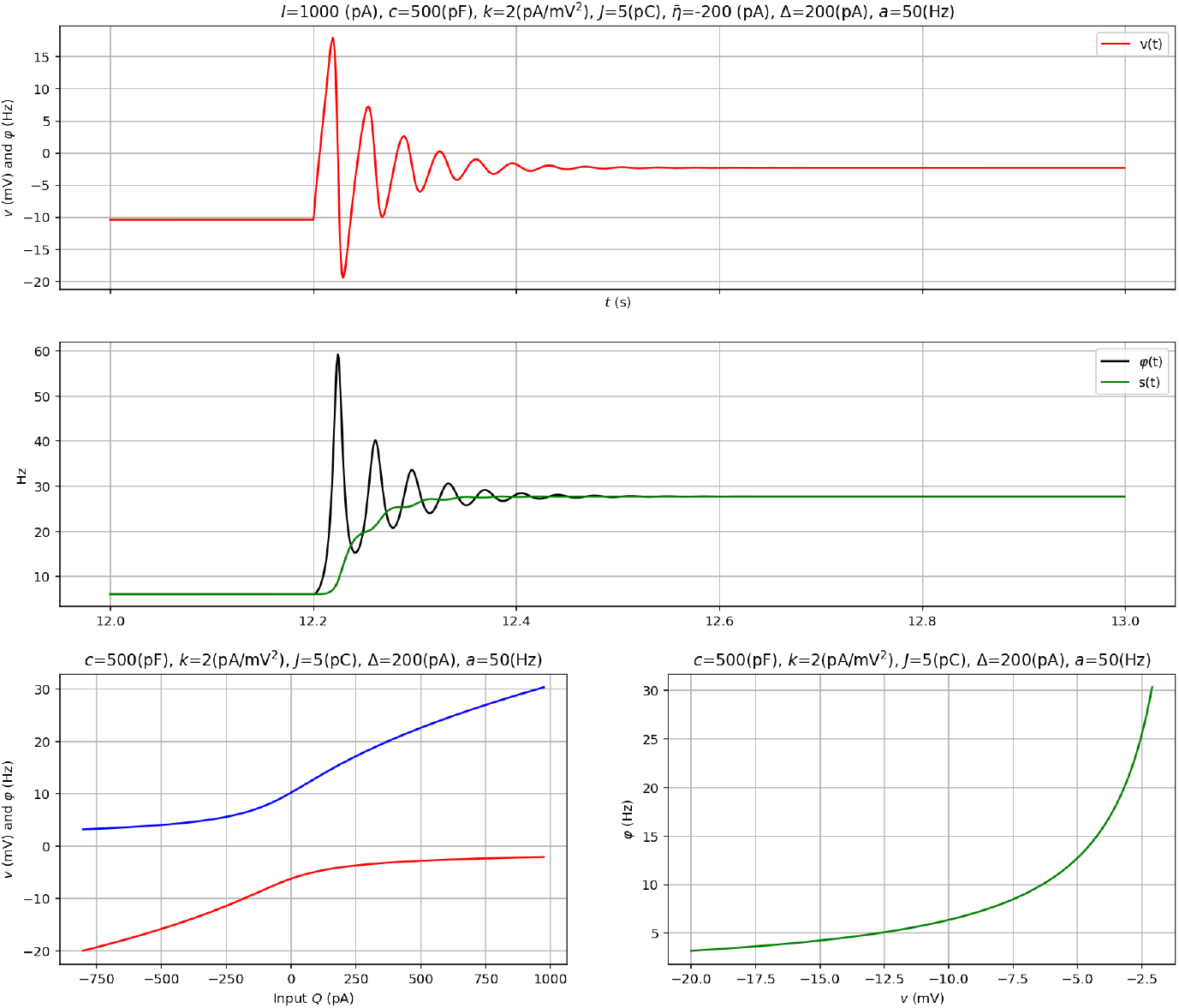
Sample dynamics and response/W2P function for selected parameters.

**Fig. SI-17.**
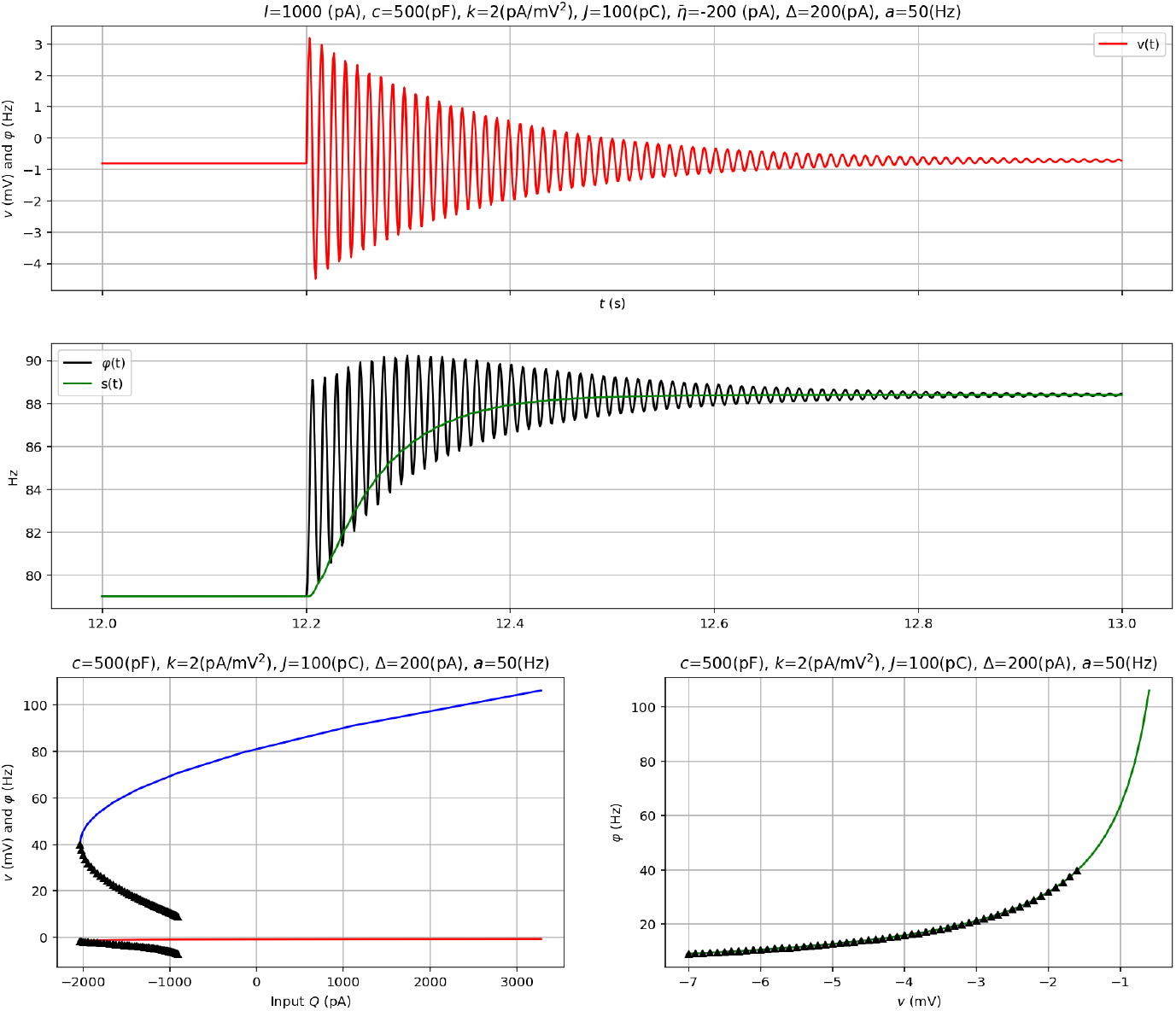
Sample dynamics and response/W2P function for selected parameters.

**Fig. SI-18.**
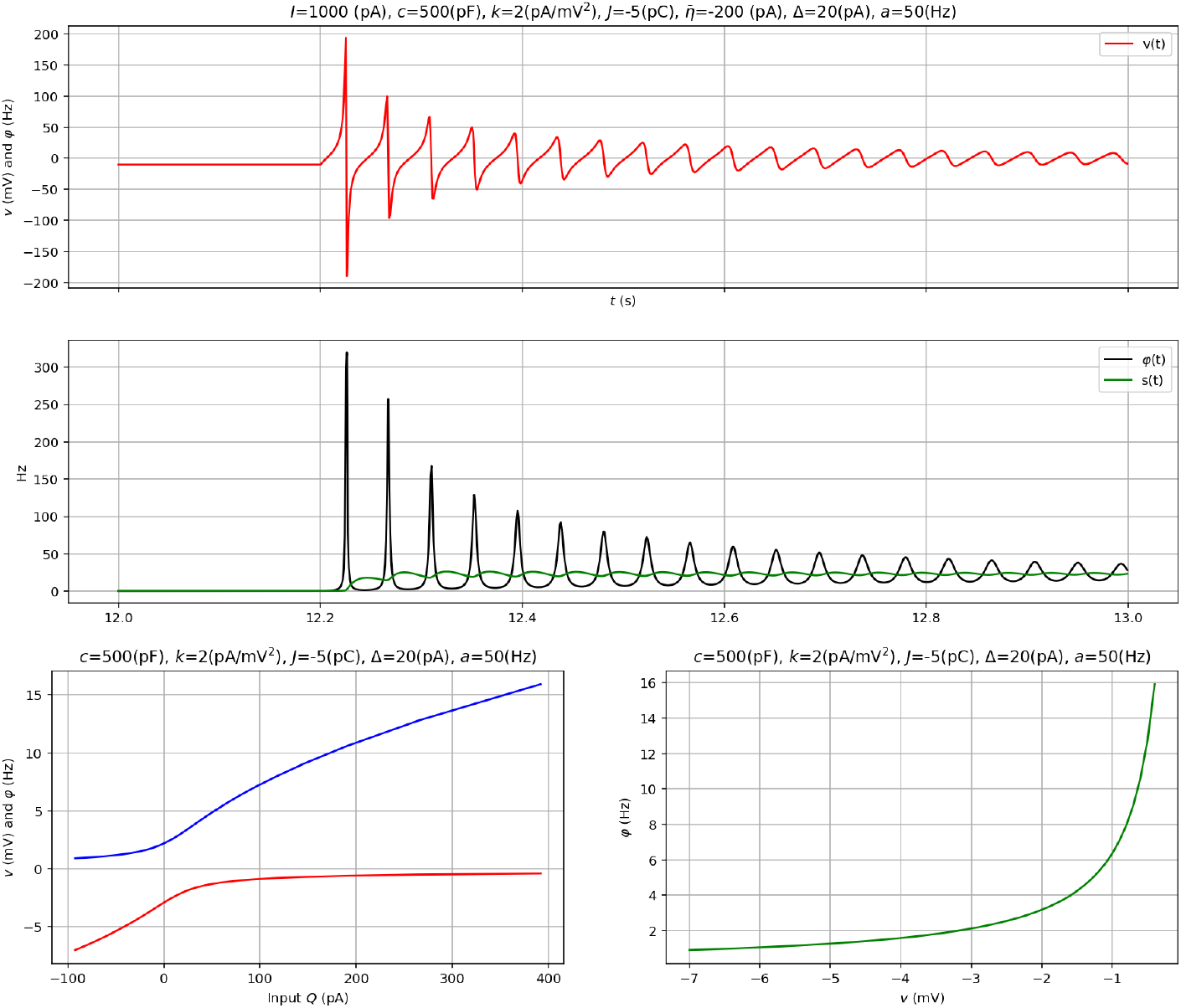
Sample dynamics and response/W2P function for selected parameters.

**Fig. SI-19.**
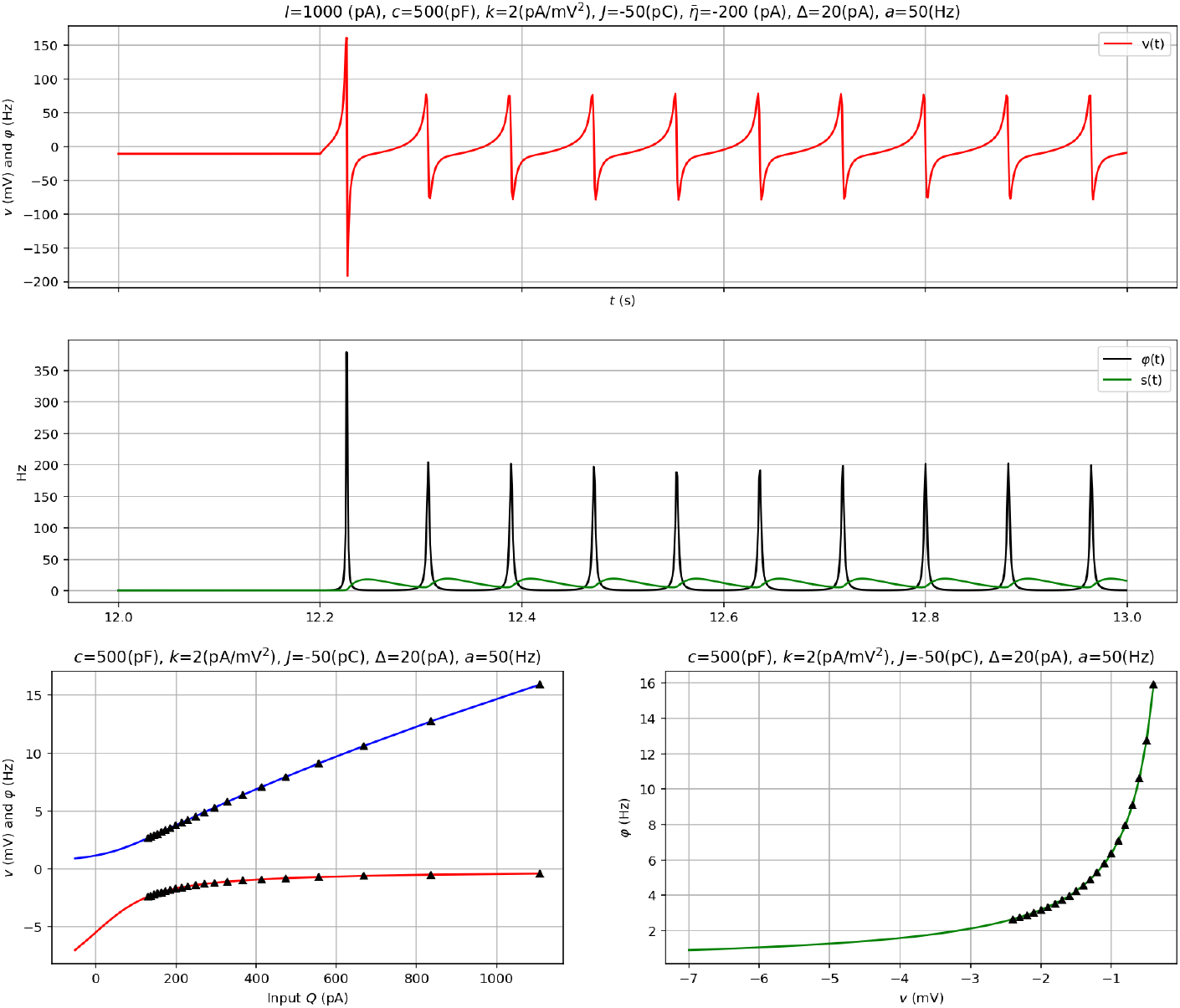
Sample dynamics and response/W2P function for selected parameters.

**Fig. SI-20.**
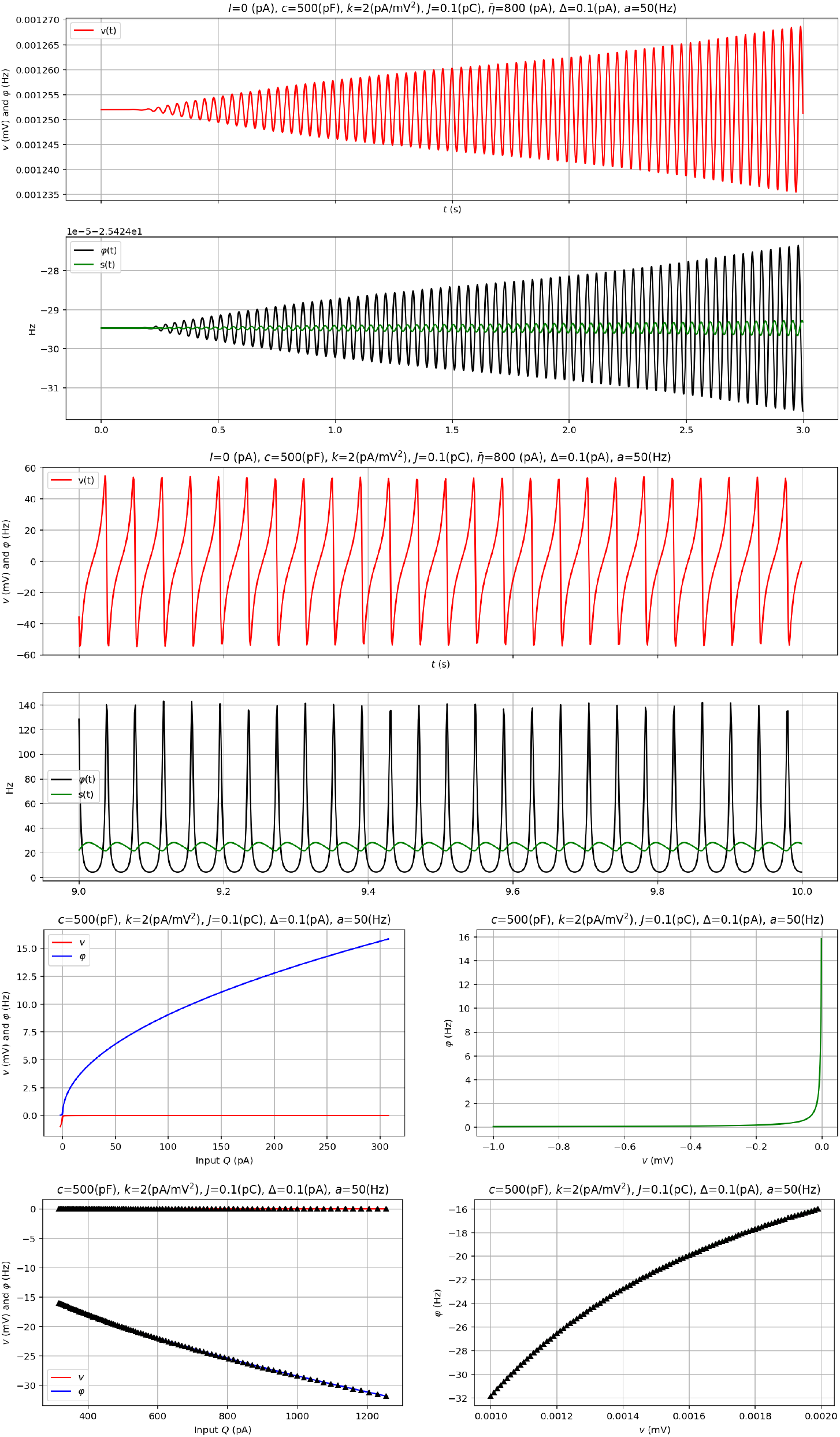
Configuration with low self-connectivity and high synchronicity at *Q* = 800. The fixed point (stable) is in a topologically disconnected space (negative firing rate, positive potential). An initial condition in the right space converges to fixed point (top), an initial condition on the other side orbits, reproducing QIF dynamics. W2P function common to both is provided in the bottom plots. Fixed points with positive firing rate occur are stable but require lower inputs.

### H. Extension to gap junctions

We can extend the analysis to the model discussed in (16), including gap junctions and asymmetric spikes. Let *g* denote conductance (units of 1*/*Ohm or [A/V]).

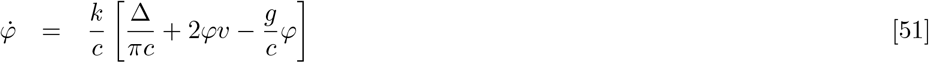

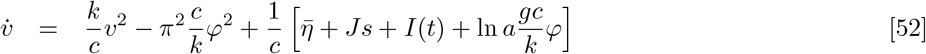

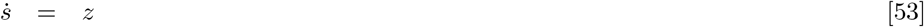

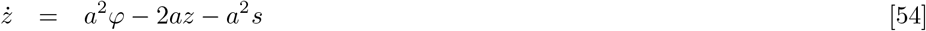

### I. Dimensional reduction of single population model

The extended equations of the single population model are ^∗^

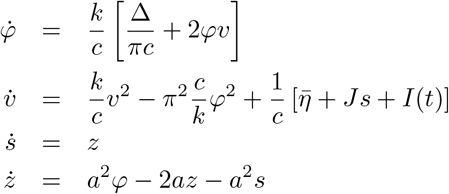

with units [*r, s*]=Hz, [*J*]=C, [*v*]=V, [*k*]=A/V^2^, [*c*]=C/V=A/(V Hz), 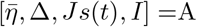, and [*k/c*]=Hz/V, [*a*] = Hz. Some natural units in NMM2 are

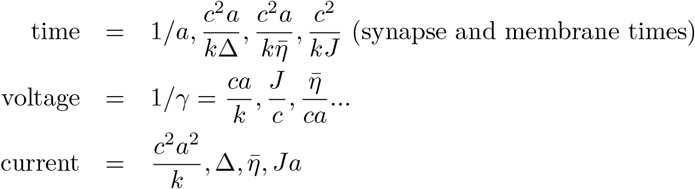

The parameters in these equations are 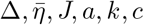 and the dimensions in the problem are *T, V, C*. The associated dimensional matrix is

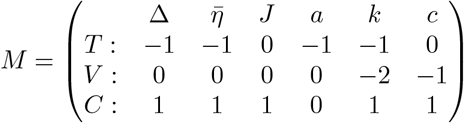

The fundamental dimensionless parameters (zero modes of the dimensional matrix) are

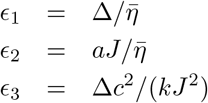

All dimensional quantities can be expressed as the product of an arbitrary one times a function of the dimensionless parameters. E.g., all frequencies can be expressed as *f* = *a F* (*ϵ*_1_, *ϵ*_2_, *ϵ*_3_).

Let the a time variable be *τ* = *at*, with 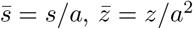, and 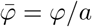. Then, using a prime to denote derivative w.r.t to *τ*,

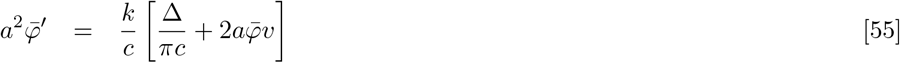

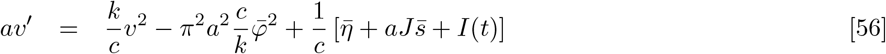

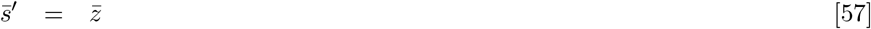

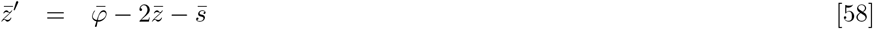

Define *γ* = *k/*(*ca*), with units [*γ*]= 1 /V and the dimensionless variable 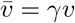, then

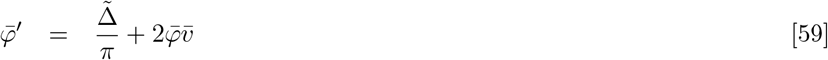

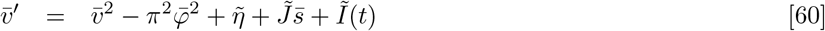

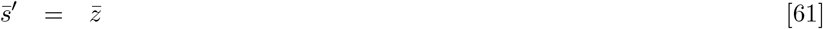

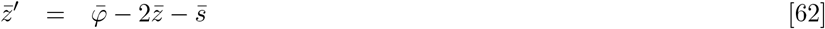

with the dimensionless current parameters

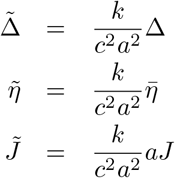

#### Stability analysis

We analyze here the fixed points and their stability of the NMM2 single population model.We start from the equations

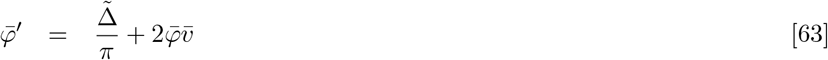

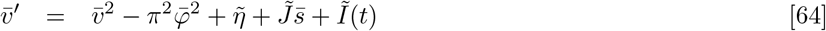

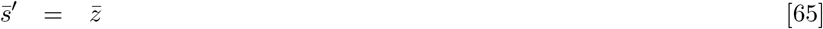

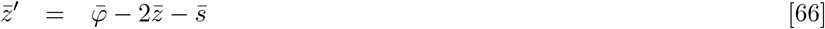

Fixed points, as described in the methods section, Equation 33, are determined by

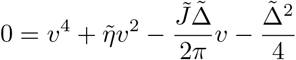

The Jacobian is

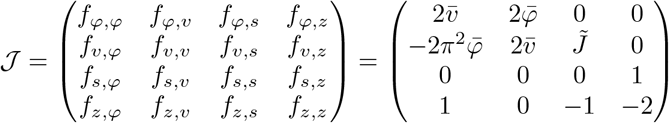

or, using the equilibrium condition 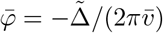,

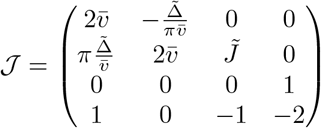

### J. Response to AC stimulation

**Fig. SI-21.**
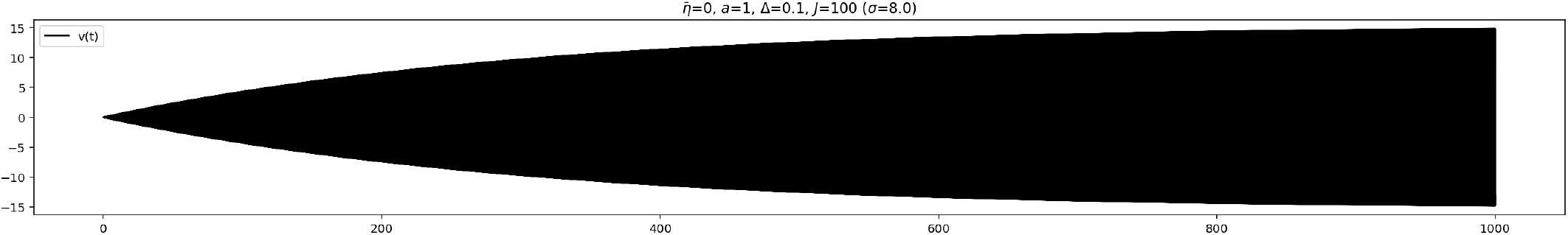
Sample time-series of *v* with resonant frequency (around 10 Hz) stimulation starting from fixed points. The rise time to steady state can be very slow and is proportional to the amplification factor of the amplitude.

**Fig. SI-22.**
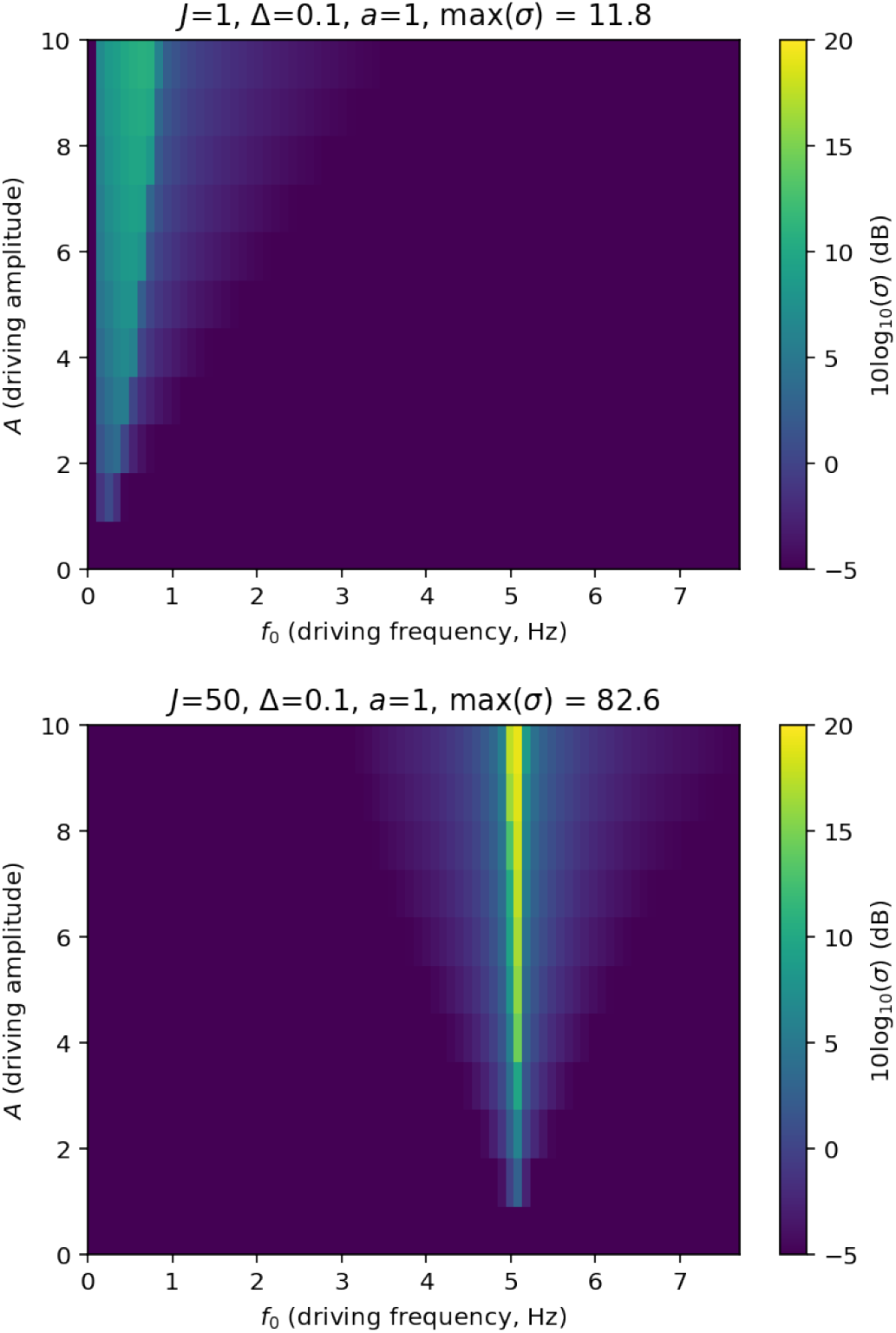
Sample Arnold tongues for AC stimulation with a weak E-field.

**Fig. SI-23.**
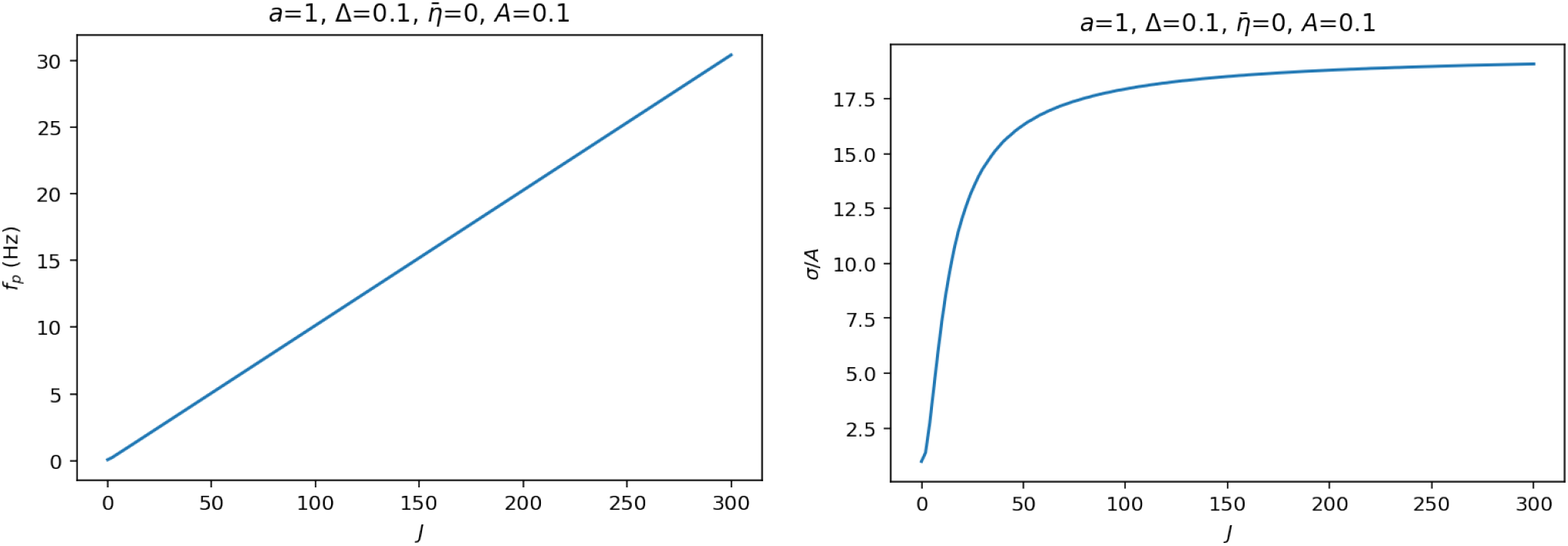
Frequency and amplitude as a function of *J* (bottom).

**Fig. SI-24.**
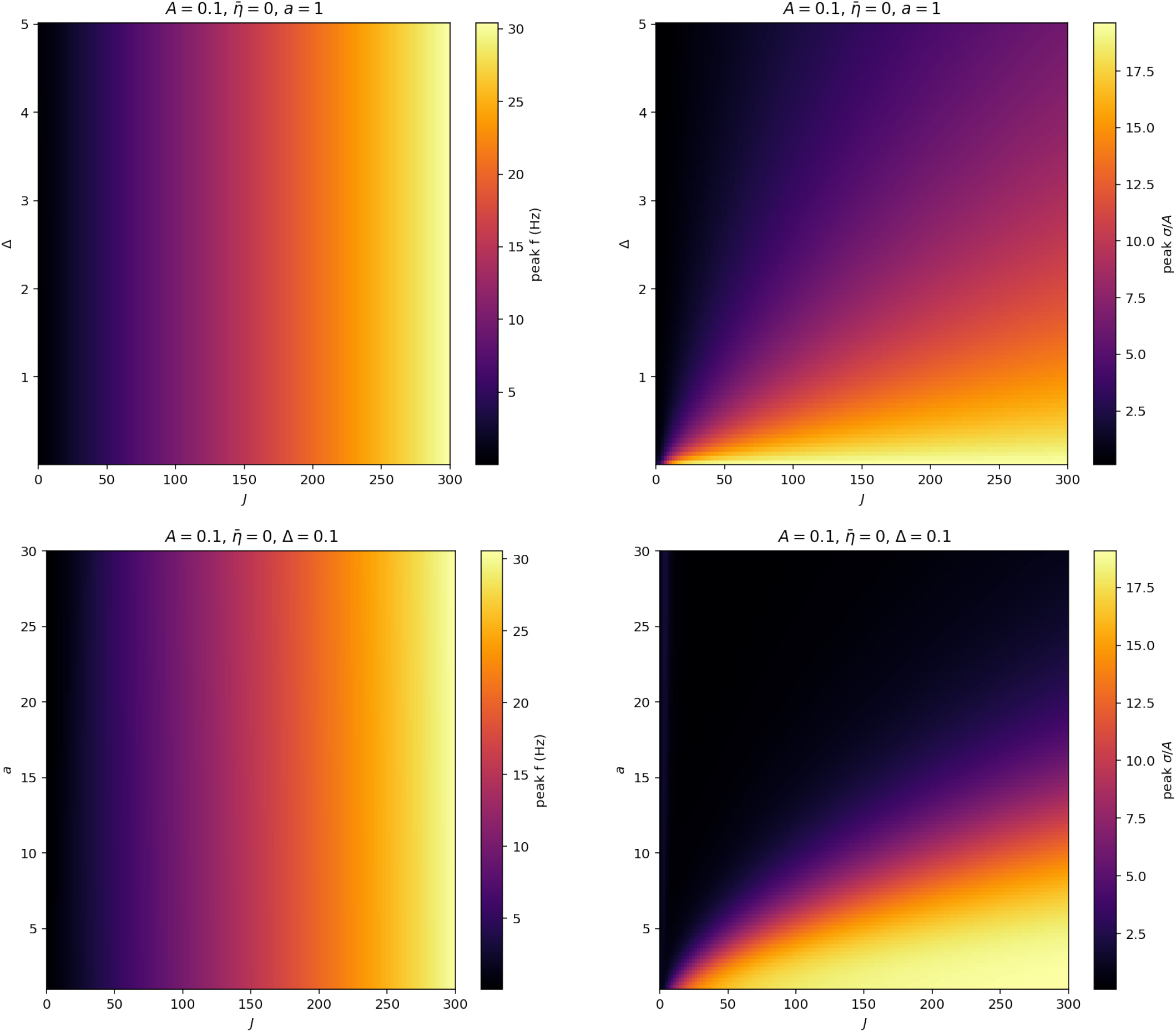
Response frequency (left) amplitude (right) as a function of *J* and Δ (top) or *J* and *a* (bottom).

### K. Slow synapse/slow input limit and mapping of NMM2 to NMM parameters

What happens in the limit of “frozen synapses”, i.e., *a* = 0^+^? As we have seen after nondimensionalization of these equations, if the dimensionless currents are large (e.g., *a*→ 0), the first two equations have large derivatives (moving fast compared to the second two) and can be evaluated at their equilibrium point (assuming they have a stable one, as we know they do from MPR), and we are in the regime of slow synapse time (compared to membrane time). The equations become — if also the input current is static or slowly varying so that the *v, φ* subsystem has time to reach equilibrium,

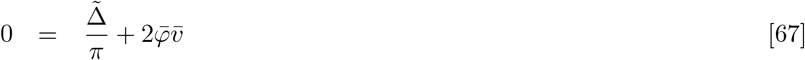

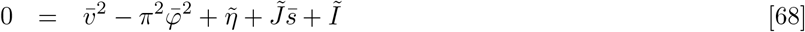

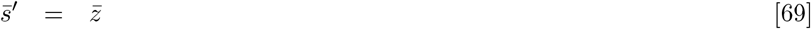

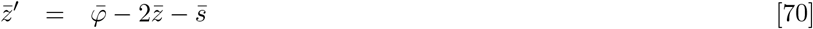

with the dimensionless current parameters

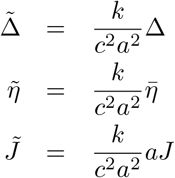

We are thus led to a traditional NMM formulation with a fixed W2P function. The details of this follow.

But we should note that it is important that the input current *I* be static or very slowly varying. The first two equations will then have time to stabilize going to a fixed point.

Assuming the above, we would like to obtain from this the wave2pulse (W2P) relation: firing rate from membrane potential or current perturbation from baseline, that is, a relation of the form (we drop all bars and tildes for simplicity)

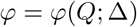

with the total input *Q* = *η* + *Js* + *I*(*t*).

Then the differential equations become

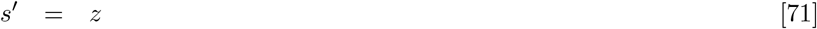

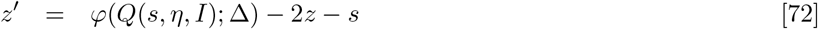

Note that in this form, the NMM2 equations (in the limit of slow synapses compared to other timescales) look very much like NMM equations in a self-coupled population. The main difference is the fact that W2P is now replaced by a different function, and that the fundamental dynamical variable is now synaptic current rather than the synaptic potential alteration *u*.

This is actually advantageous, because to connect with physical measurements we are interested in synaptic currents, and, in fact, in the NMM formalism we use a linear relation to transform *u* into *s*.

The main issue with the NMM2 formulation is the fact that the WP2 function does not saturate, a fact related to the lack of refractory period in the QIF equations (the QIF neuron can increase its firing rate to infinity as a function of input current).

To get closer to the NMM formulation, we write

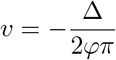

and input this into the second equation above,

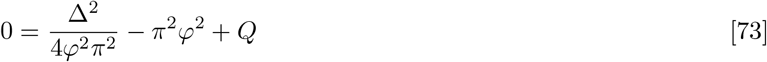

or

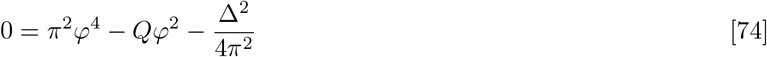

or

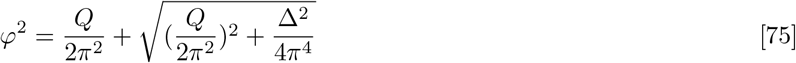

or (since *φ* > 0),

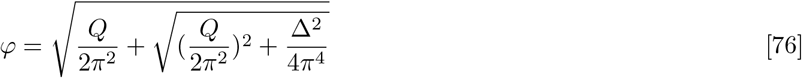

#### Fixed point at I = 0

We start from *φ*\* = *s*\* and plug this into equation 73 to get

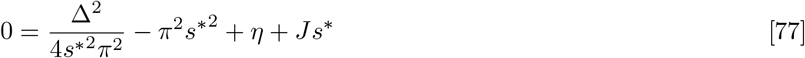

#### Derivative

We can compute the derivative

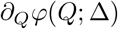

differentiating equation 75 to get

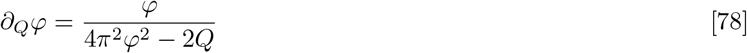

and evaluating at the fixed point

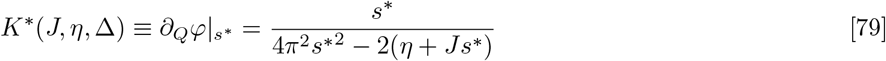

We can use equation 75 above with *φ*\* = *s*\* to get

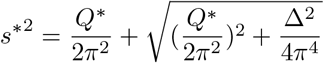

From this we see that 4*πs*^*2^ > 2*Q*^*2^, so *K*\* ≥ 0. More explicitly,

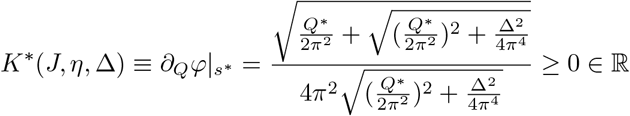

Since *Q*\* = *η* + *Js*\* + *I*, from equatio 78 we see that the derivative *φ*′(*s*) at the fixed point is

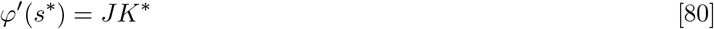

and therefore has the same sign as *J*.

#### Linearized equations

We can then linearize these equations about their equilibrium points (*s*\*, *z* = 0), *φ*(*s*\*) − *s*\* = 0 and for small *I*, and defining *t* = *s* − *s*\* write

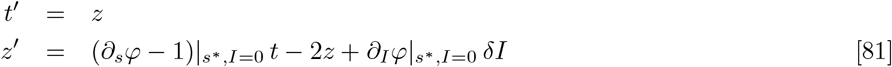

or

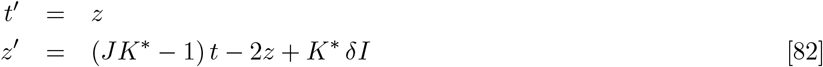

In second order form

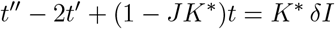

The eigenvalues of this ODEs are (recall *K*\* ≥ 0),

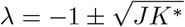

We note here that for *J* > 0 the eigenvalues are real (no resonance possible), but recall that the slow synapse approximation is applicable only if the dynamic external input to the population is slowly varying w.r.t. QIF dynamics.

#### Resonance for small perturbations

If *J* < 0, The eigenvalues of this ODEs are

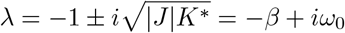

The resonance peak over amplitude is of the form

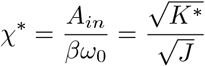

Plots for fixed point, *K*\* and *χ*\* as a function of *J* are provided in Figure SI-25.

#### Relation to NMM sigmoid

The NMM sigmoid is given by

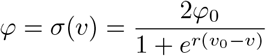

How similar are the two transfer/W2P functions? Some examples of fitting are provided in Figure SI-25.

**Fig. SI-25.**
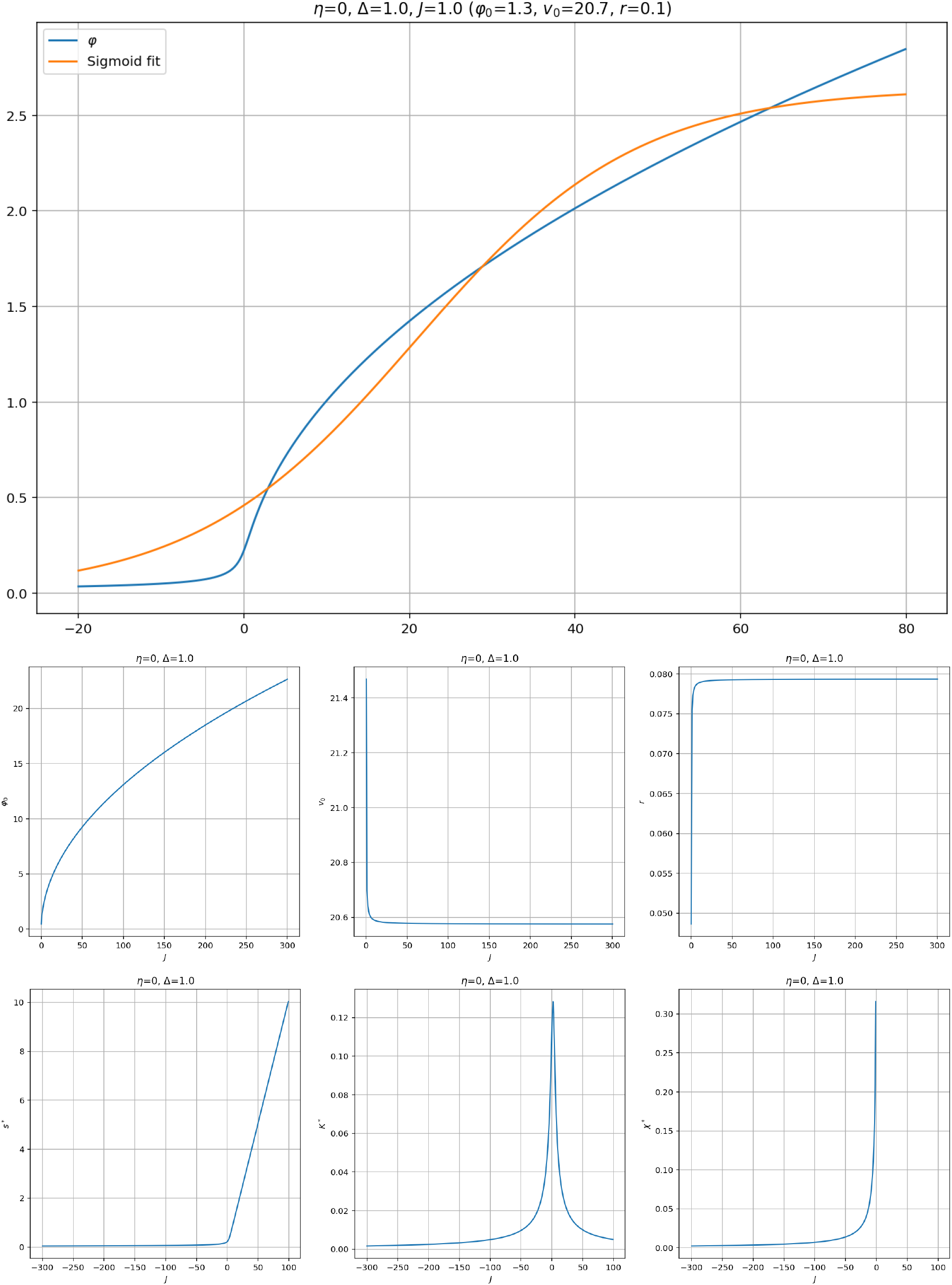
NMM2 slow limit. Fitting of a sigmoid to the transfer function for some parameters (top) and fit sigmoid parameters for a range of *J* values (middle). The most important change is for the firing rate. Bottom: fixed point (*s*\* = *φ*\*), derivative *K*\* and resonant peak for various values of *J* (there is no resonance for positive *J* values).

### L. Slow synapse/fast input limit

In the limit of slow synapses (*a* = 0^+^) but fast input, we have that we need to treat the synaptic input as a constant *s*\* (it does not change at all compared to the the other time scales and we take it as frozen in time). The equations become (we drop bars and tildes),

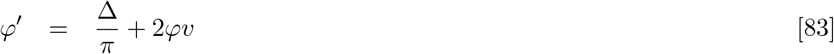

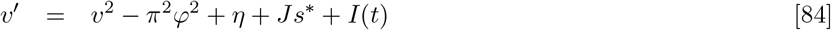

The Jacobian to be evaluated at the fixed point *v*\*, *φ*\* is

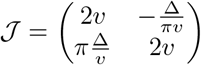

and the eigenvalues are

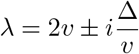

The resonant peak in the linear regime is 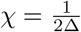.

### M. Formal relation between NMM2* and NMM

The formal relationship between NMM2 and NMM occurs in the limit of slow *Q* = *η* + ∑*Js* + *I*(*t*), that is, when all the inputs to the population are slow compared to MPR/QIF dynamics (we can call this the NMM2* regime). In that case we can formally related the two theories through their transfer functions as summarized in Figure SI-26.

**Fig. SI-26.**
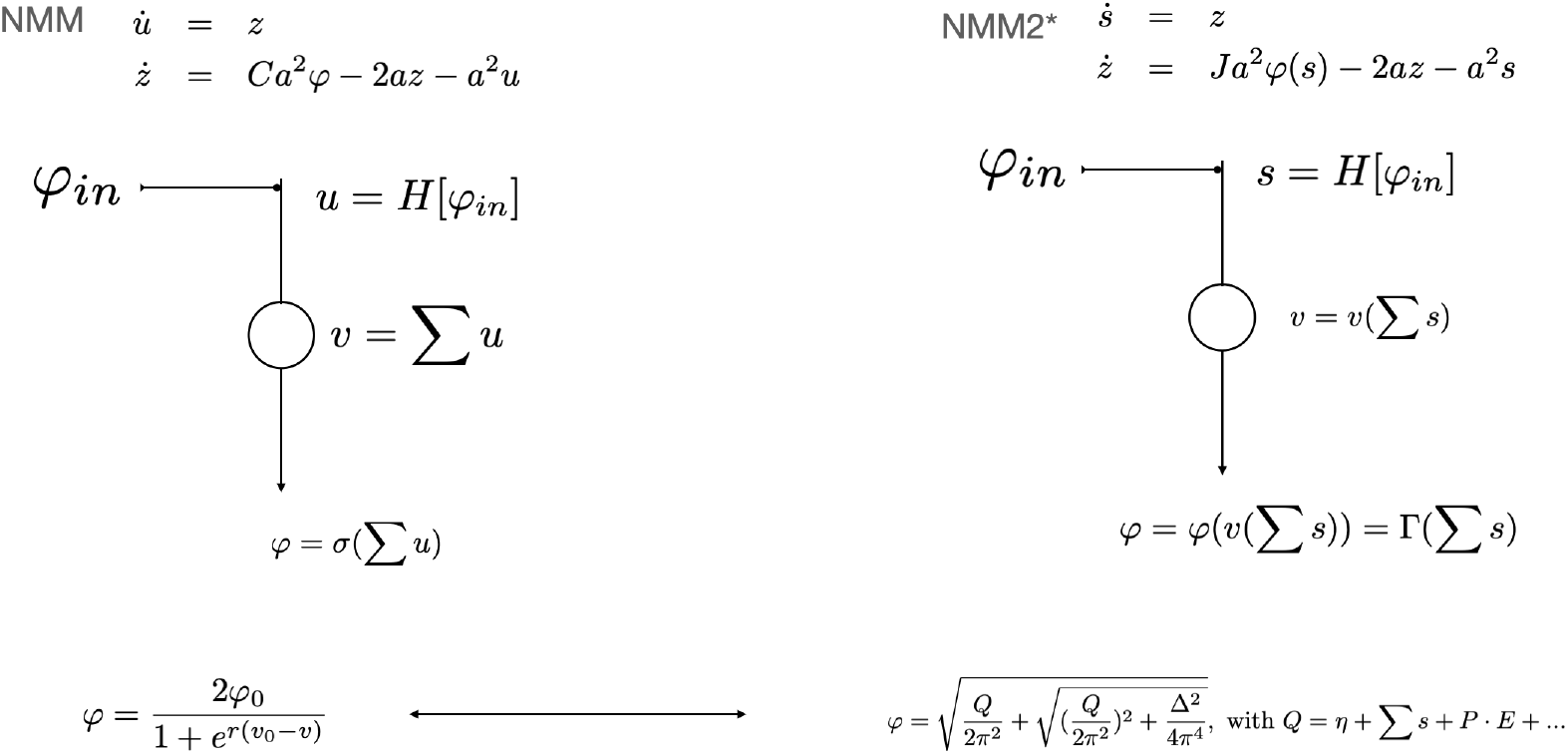
Relationship of NMM2* and NMM.

### N. Further notes on the relation with the Devalle et al formulation (17)

Devalle et al (17) start from the QIF equation

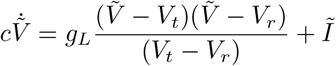

Here we have the proper physical units: [*V*] = V, [*Ĩ*] = A, [*gL*] = A/V, etc.

Aside: Note that the dynamic quantity

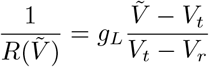

has units of conductance (Ω^− 1^), and with it we can rewrite the differential equation in an intuitive form (RC circuit),

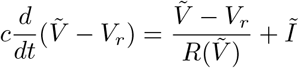

The time “constant” associated to this RC circuit would be

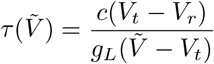

#### Connection with our notation

We define

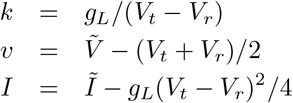

then we obtain

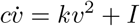

This is the formulation we start from here. Here we preserve units of all the variables, with [*v*]=V, [*I*] =A, [*k*]=A/V^2^, [*c*]=C/V=A/(V Hz), and [*k/c*]=Hz/V.

Again, we can refer here to a dynamical time scale if we want to think in terms of an RC circuit,

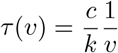

#### Devalle time scale

In order to get a time scale in the system we need to use other constants — the threshold or rest voltage. Here we use 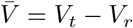 as is the case in Devalle.

Dimensional analysis shows that 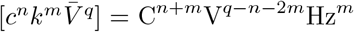, and using *m* = 1, *n* = − 1, *q* = 1 we find the desired timescale,

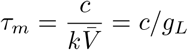

Devalle also rescale 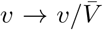 to and 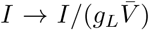 to end up with the almost adimensional equation (adimensional except for time) 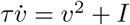. Rescaling the time variable then leads to 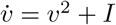.

#### Other time scales

More generally, the dimensional analysis can include the input *I*:

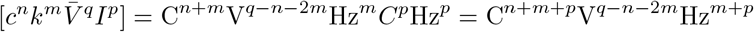

Setting *m* + *p* = − 1, or *p* = − 1 − *m*

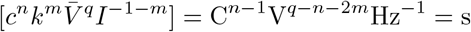

thus we need *n* = 1, *q* − 1 − 2*m* = 0, or

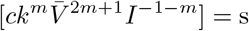

E.g., setting *m* = 0, we get the time scale

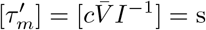

Setting *m* = − 1*/*2,

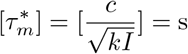

What does this all mean? That we can rewrite the DE in multiple ways. E.g., start from

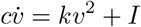

And multiply it by 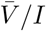,

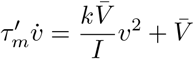

We can now rescale *v* = *αv*^*′*^,

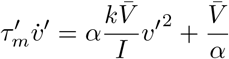

or with 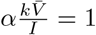

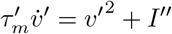

With 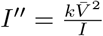. So, as we can see, there are different transformations that lead to the same form 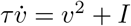, but with different time scales. This may be relevant for fast-time slow-time analysis.

From the point of view of connecting to physiology, though, the version that makes most sense to work with is the Devalle one: time constant of the membrane (which is what is related to PSC measurements) is not related to input current.

### O. Units systems in QIF, MPR and NMM2

#### QIF

Consider a population of fully and uniformly connected quadratic integrate and fire (QIF) neurons indexed by *j* = 1, …, *N*. To make a direct connection with experimental work, we start from the equation for the membrane potential of a single neuron *V*^*j*^ in a population of interest as in (17),

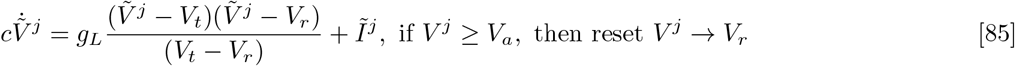

where *V*_*r*_ and *V*_*t*_ represent the resting potential and threshold of the neuron (in Volts), and *V*_*a*_ is a limit reset potential (apex), 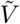 is the membrane voltage, *Ĩ* the input current, *c* the membrane capacitance, *g*_*L*_ is the leak conductance, with units 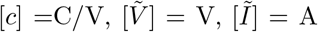 and [*g*_*L*_] = A/V. In this equation, the total input current to neuron *j* is 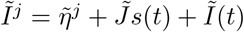 and includes a quenched (constant) noise input component 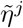 with mean 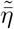 and variance 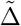, the input from other neurons *s*(*t*) per connection received (the mean synaptic activation) with uniform coupling 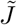 (with units of charge) and a common input *Ĩ*(*t*). The common input *Ĩ*(*t*) can represent both a common external current input or the effect of an electric field.

#### Natural units

To simplify the analysis (completing the square), we define

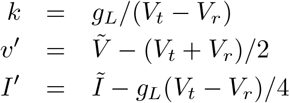

which, through *Ĩ*, also affects 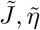 and 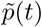, to obtain

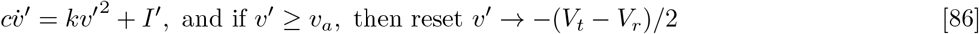

with units [*k*]= A/V^2^ and voltage and current in proper units (V and A, respectively). Defining *v* = *v′/*(*V*_*t*_ − *V*_*r*_) and *I* = *I′/*(*g*_*L*_(*V*_*t*_ − *V*_*r*_)) results in

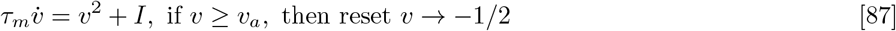

with *τ*_*m*_ = *c/g*_*L*_ and *v, I, η* and *p*(*t*) dimensionless variables, and *J* with units of time (as *s* is in Hz). If we work in time units defined by the timescale *τ*_*m*_, the equation becomes

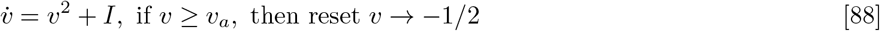

What we have done by the above transformations is essentially work in natural units of time (*τ*_*m*_ = *c/g*_*L*_), voltage (*V*_*t*_ − *V*_*r*_) and current (*g*_*L*_(*V*_*t*_ − *V*_*r*_)). It is important to keep in mind these changes of variables when dealing with multiple interacting populations involving different parameters. The coupling parameters across populations as well with electric field are affected by the above transformations.

Thus, the above operations are essentially equivalent to working in a system of units where

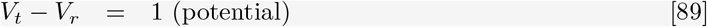

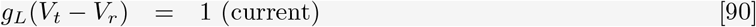

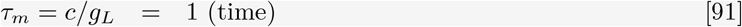

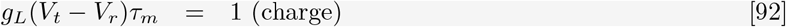

#### MPR

Following the derivation of the mean-field equations in (15) we get

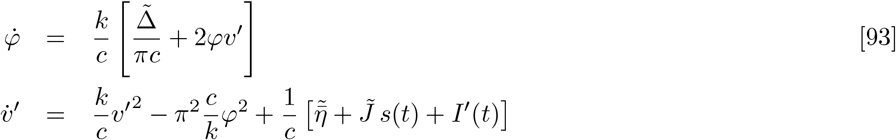

with units 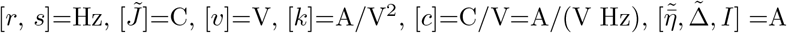 and [*k/c*]=Hz/V. To close these equations, we write 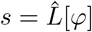 for some differential operator representing a causal, linear filter representing synaptic dynamics. The cases of first and second order filtering are discussed below. In the limit of instantaneous synaptic transmission, *τ* →0 or *a*_*τ*_ (*t*− *t′*) →*δ*(*t*− *t′*), which implies *s*→ *φ*, we obtain the closed, simple set of equations for the single population model analyzed in (15).

In the reduced version with the variables and units as in Equation (87) — where *V*_*t*_ − *V*_*r*_ = 1 (potential) and *g*_*L*_(*V*_*t*_ − *V*_*r*_) = 1 (current) —, the MPR equations become (17)

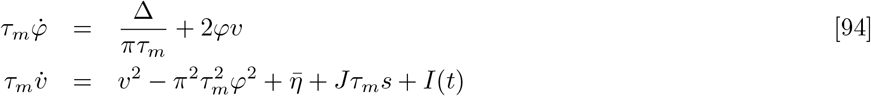

The units are: voltage *v* is dimensionless, the rates *φ* and *s* have units of frequency (Hz), *τ*_*m*_ = *c/g*_*L*_ has units of time (seconds), and the charge 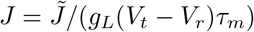 and current 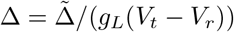 dimensionless.

We can define time units measured by *τ*_*m*_ (so that in the new units *τ*_*m*_ = 1) by using a new times variable 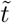, with 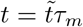. We use tildes to denote rates and their derivatives in the new time units thorough multiplication by *τ*_*m*_. Then the equations transform to (with tilde denoting rates measured in these units)

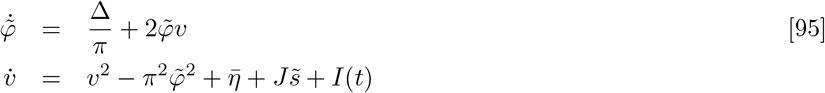

which is the formulation in (15). All variables and parameters are dimensionless and expressed in the natural units of the problem, where *τ*_*m*_ = *c/g*_*L*_ = 1 (time), *V*_*t*_ − *V*_*r*_ = 1 (voltage), *g*_*L*_(*V*_*t*_ − *V*_*r*_)*τ*_*m*_ = 1 (charge) and hence *g*_*L*_(*V*_*t*_−*V*_*r*_) = 1 (current).

#### Formulation for the single and multiple population cases

The complete set of equations for a single population becomes

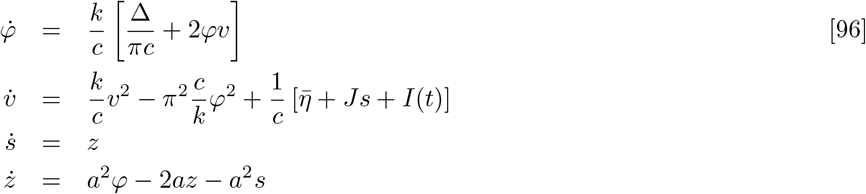

with the input collecting external input and the influence of an external field, *I*(*t*) = *p*(*t*) + *P·E*(*t*).

The equations for the single population can be reduced to the simplified form in Equation (94) with dimensionless voltage *v* and current *I* that brings to light the two main timescales in the problem,

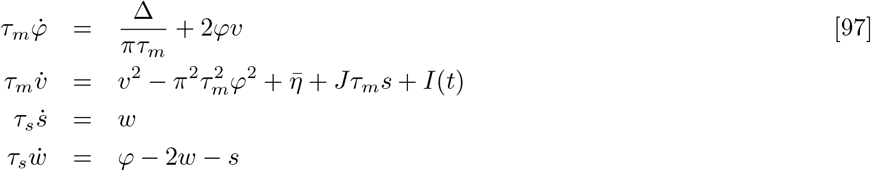

with *τ*_*s*_ = 1*/a* and and *w* = *τ*_*s*_*z*. Here *s* and *w* have units of Hz, while *v* is dimensionless, *φ* and *s* are in Hz, *τ*_*m*_ = *c/g*_*L*_ has units of time (seconds), 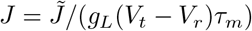 is the dimensionless charge, and 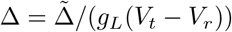 the dimensionless current.

As before, can transform the above equations to dimensionless units by using *τ*_*m*_ as the time unit (*τ*_*m*_ = 1),

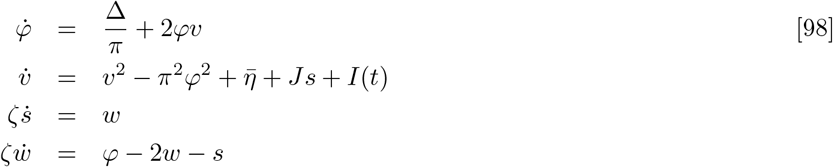

All variables are dimensionless and 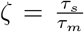 the synaptic time constant in membrane time units. The first two equations are again the formulation in (15).

Equivalently, we can multiply the last equation by *a*^2^, with *a* = *ζ*^− 1^ = *a*_*s*_*/a*_*m*_ (i.e, the synaptic rate in the *τ*_*m*_ unit system),

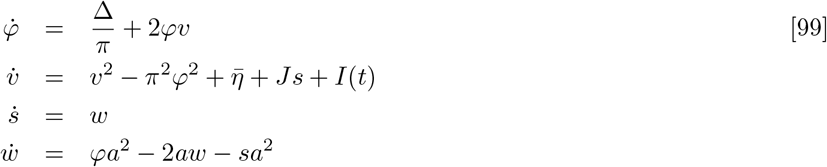

In summary, we can write the NMM2 equations in units defined by *τ*_*m*_ = 1 (time), *V*_*t*_ − *V*_*r*_ = 1 (voltage) and *g*_*L*_(*V*_*t*_ − *V*_*r*_) = 1 (current), *g*_*L*_(*V*_*t*_ − *V*_*r*_)*τ*_*m*_ = 1 (charge) in this simplified form.

#### Multiple populations

The single population case in Equation (96) is readily generalized for interacting populations. Letting *p* denote the population index and *pq* denote a synapse from population *p* to *q*, the uniform input received by population *p* by *I*_*p*_(*t*) = *p*_*p*_(*t*) + *P*_*p*_ · *E*_*p*_(*t*), the equations for interacting populations become

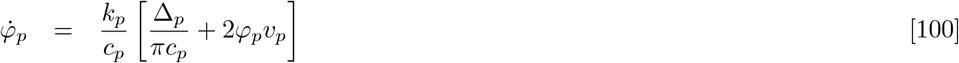

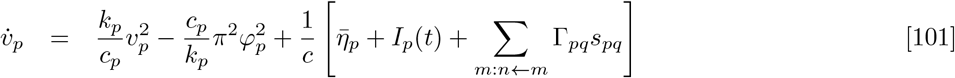

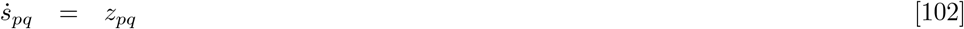

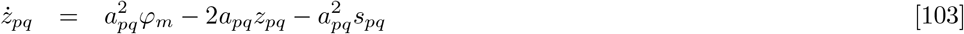

Figure 1 provides a diagram of the self-coupled population and multiple population cases.

Finally, we can work with the form in Equations (97) by defining dimensionless voltage variables independently in each population if need be (i.e., if *V*_*t*_ − *V*_*r*_ and *g*_*L*_(*V*_*t*_ − *V*_*r*_) differ among them) and taking care of the coupling between them and with external electric fields. Each population has its own unit system, so to speak.

## Footnotes

∗ One may be tempted to replace the last equation by *z* = *a*^2^ *βφ* − 2*az* − *a*^2^ *s*, but this is redundant. The change of variables 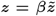 and 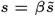 then lead to 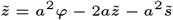, and the *β* factor pops up in the second NMM2 equation, changing *J* → *βJ*. This is also clear from the point of view of the *L* operator.

